# Loss of Activating Transcription Factor 3 prevents KRAS-mediated pancreatic cancer

**DOI:** 10.1101/2020.03.27.011601

**Authors:** Nawab Azizi, Jelena Toma, Mickenzie Martin, Muhammad Faran Khalid, Nina Steele, Jiaqi Shi, Marina Pasca di Magliano, Christopher L. Pin

**Affiliations:** Children’s Health Research Institute, London, Ontario, N6C 2V5, Canada; Department of Paediatrics, University of Western Ontario, London, Ontario, N6A 3K7, Canada; Department of Physiology and Pharmacology, University of Western Ontario, London, Ontario, N6A 3K7, Canada; Department of Oncology, University of Western Ontario, London, Ontario, N6A 3K7, Canada; Department of Biology, University of Western Ontario, London, Ontario, N6A 3K7, Canada; Dept. of Pathology, University of Michigan, Ann Arbor, Michigan, USA; Dept. of Surgery, University of Michigan, Ann Arbor, Michigan, USA

**Keywords:** Unfolded Protein Response, ER stress response, pancreatic ductal adenocarcinoma, acinar to ductal metaplasia

## Abstract

The unfolded protein response (UPR) is activated in pancreatic pathologies and suggested as a target for therapeutic intervention. In this study, wxe examined Activating Transcription Factor 3 (ATF3), a mediator of the UPR which promotes acinar-to-ductal metaplasia (ADM) in response to pancreatic injury. Since ADM is an initial step in the progression to pancreatic ductal adenocarcinoma (PDAC), we hypothesized ATF3 is required for initiation and progression of PDAC. We generated mice carrying a germ line mutation of *Atf3* (*Atf3*^*-/-*^) combined with acinar-specific induction of oncogenic KRAS (*Ptf1a*^*creERT/+*^*Kras*^*LSL-G12D*^). *Atf3*^*-/-*^ mice with (termed *APK*) and without KRAS^G12D^ were exposed to cerulein-induced pancreatitis. In response to recurrent pancreatitis, *Atf3*^*-/-*^ mice showed decreased ADM and enhanced regeneration based on morphological and biochemical analysis. Similarly, an absence of ATF3 reduced spontaneous pancreatic intraepithelial neoplasia formation and PDAC in *Ptf1a*^*creERT/+*^*Kras*^*LSL-G12D*^ mice. In response to injury, KRAS^G12D^ bipassed the requirement for ATF3 with a dramatic loss in acinar tissue and PanIN formation observed regardless of ATF3 status. However, unlike *Ptf1a*^*creERT/+*^*Kras*^*LSL-G12D*^ mice, *APK* mice exhibited a cachexia-like phenotype, did not progress through to PDAC, and showed altered pancreatic fibrosis and immune cell infiltration. These findings suggest a complex, multifaceted role for ATF3 in pancreatic cancer pathology.

## Introduction

The unfolded protein response (UPR) is a critical pathway protecting cells from a variety of harmful stresses that promote improper protein folding or processing [1, 2]. In response to accumulation of misfolded proteins, the UPR is activated to reduce protein load through a general decrease in protein translation, activation of ER-associated degradation, and increased expression of protein chaperones [1-3]. Over the last decade, the UPR has been implicated in a variety of human pathologies, including breast [4], lung [5] and colorectal cancer [6, 7]. In the pancreas, the UPR plays important roles in both physiological and pathological processes. More recently, the UPR has been suggested as a mediator and potential therapeutic target for pancreatic ductal adenocarcinoma (PDAC; [8, 9]. PDAC is currently the third leading cause of cancer-related deaths, with a dismal 5-year survival rate of ∼9%. While constitutive activation of KRAS is an initiating event in PDAC, targeting this pathway has been futile to date, suggesting alternative targets need to be identified. Studies with pancreatic cancer cell lines indicate that targeting the UPR may be beneficial [8, 10]. However, the mechanism(s) of action of the UPR in cancer initiation and progression have been elusive. This is likely due to the numerous levels of regulation that exist for the UPR both before and after activation.

Within acinar cells, the UPR plays a pivotal role in normal physiology, preventing accumulation of misfolded proteins in the ER lumen and maintaining protein homeostasis [11]. The UPR consists of three signalling branches - PERK ([PKR]-like ER kinase), IRE1 (inositol requiring enzyme 1) and activating transcription factor 6 (ATF6; [12]. These ER-membrane associated proteins are triggered through dissociation of GRP78/BiP in response to accumulation of misfolded protein [13]. Activation of PERK leads to phosphorylation of eIF2α which then leads to global inhibition of mRNA translation [14]. An important mediator of PERK signaling is ATF4, which is transcribed but in the absence of stress, but not translated as a functional protein [15]. Upon stress, *Atf4* avoids translational inhibition, is translated from an alternative promoter, and activates genes that regulate protein folding, degradation and cell survival [16, 17]. PERK signaling also activates *Atf3* gene expression [18].

Likely due to the high protein turnover rate in acinar cells [19], deletion of genes encoding several important mediators of the UPR, including PERK [20], IRE1 [19] and ATF4 [20], all result in pancreatic pathology highlighting their importance to acinar cell physiology. To understand the pathological requirement of the UPR, we have focused on ATF3, which is not expressed in the pancreas until induced by injury, such as observed in pancreatitis [21]. Previous work from our laboratory and others showed ATF3 expression is rapidly induced in acinar cells by experimental forms of pancreatitis [22, 23]. In other organs or pathologies, ATF3 affects signaling pathways and cellular processes that are observed in pancreatitis and PDAC. ATF3 interacts with SMAD and MAPK signaling pathways [24] to relieve stress through DNA damage repair and cell cycle regulation [25], binds NF-κB and represses the expression of cytokines such as IL-6 and TLR-4 [26, 27], and is vital for neutrophil migration to areas of injury in lung tissue [28]. In the skin, ATF3 is critical for negatively regulating cancer-associated fibroblasts (CAFs) to prevent the excessive deposition of extracellular matrix (ECM) proteins that would otherwise promote carcinogenesis [29].

While roles for ATF3 has been identified in several cancers, these are cell and cancer-type dependent [6, 7, 30]. Conflicting roles for ATF3 have been reported for metastatic prostate cancer [31]. In breast cancer, high levels of ATF3 is associated with reduced patient survival and promotes tumour development and metastasis [32]. Alternatively, in colon and colorectal cancers, ATF3 overexpression reduces cell survival by exerting anti-tumorigenic effects [33, 34]. ATF3’s role in pancreatic cancer has not been examined to date.

Loss of the acinar cell phenotype or ADM (acinar to ductal metaplasia) is an early event in chronic pancreatitis and PDAC. Our laboratory showed ATF3 is required for ADM during acute injury [22] by activating *Sox9* (Sry-related high-mobility group box 9) and repressing *Mist1*, which maintains the mature acinar cell phenotype [35]. SOX9 and MIST1 are important regulators of ADM, required for [36] or limiting [37] PDAC progression through ADM/pancreatic intraepithelial neoplasia (PanIN) formation, respectively.

These studies suggest ATF3 may have an early role in progression of PDAC, possibly through regulation of key transcription factors involved in the ADM process. However, while recurrent or chronic forms of pancreatitis are a significant risk factor for PDAC, acute pancreatitis is a poor predictor of PDAC [38]. Therefore, in this study we investigated whether ATF3 is required for ADM in recurrent injury and whether oncogenic KRAS could override this requirement. Our findings suggest ADM can occur in KRAS-mediated PDAC, but ATF3 is required for progression and maintenance of high grade PanIN lesions and PDAC. In addition, it appears that ATF3 may affect multiple cell types involved in PDAC.

## Results

### ATF3 is required for activating the transcriptional program for ADM during recurrent pancreatic injury

To determine if ATF3 is critical for the loss of the acinar cell phenotype during recurrent pancreatic injury, congenic wild type and *Atf3*^*-/-*^ mice were subjected to rCIP for 14 days (Supplemental Figure S1A). Pancreatic tissue was collected one and seven days following cessation of cerulein to assess the extent of injury and regeneration, respectively. No gross morphological differences were observed between WT and *Atf3*^*-/-*^ mice during rCIP. Both cerulein-treated groups showed significant weight loss compared to saline treated groups (Supplemental Figure S1B), but the difference minimized after cerulein was stopped. One day following treatment, both cerulein-treated groups showed a significant decrease in pancreas/body weight ratio relative to saline groups (Figure 1A), with no difference between WT and *Atf3*^*-/-*^ mice. Both genotypes acquired a similar extent of damage (Figure 1B), including loss of acinar tissue based on amylase IHC (Figure 1C) and western blot analysis (Figure 1D) and fibrosis (Supplemental Figure S2). While morphological staining indicated similar accumulation of duct-like structures in cerulein-treated WT and *Atf3*^*-/-*^ tissue (Figure 2A), IHC for CK19 expression was less extensive in *Atf3*^*-/-*^ tissue (Figure 2B). Quantification of the extent of CK19 accumulation confirmed a reduction in the total amount of CK19+ cells in *Atf3*^*-/-*^ tissue compared to WT mice (Figure 2C).

**Figure 1.**
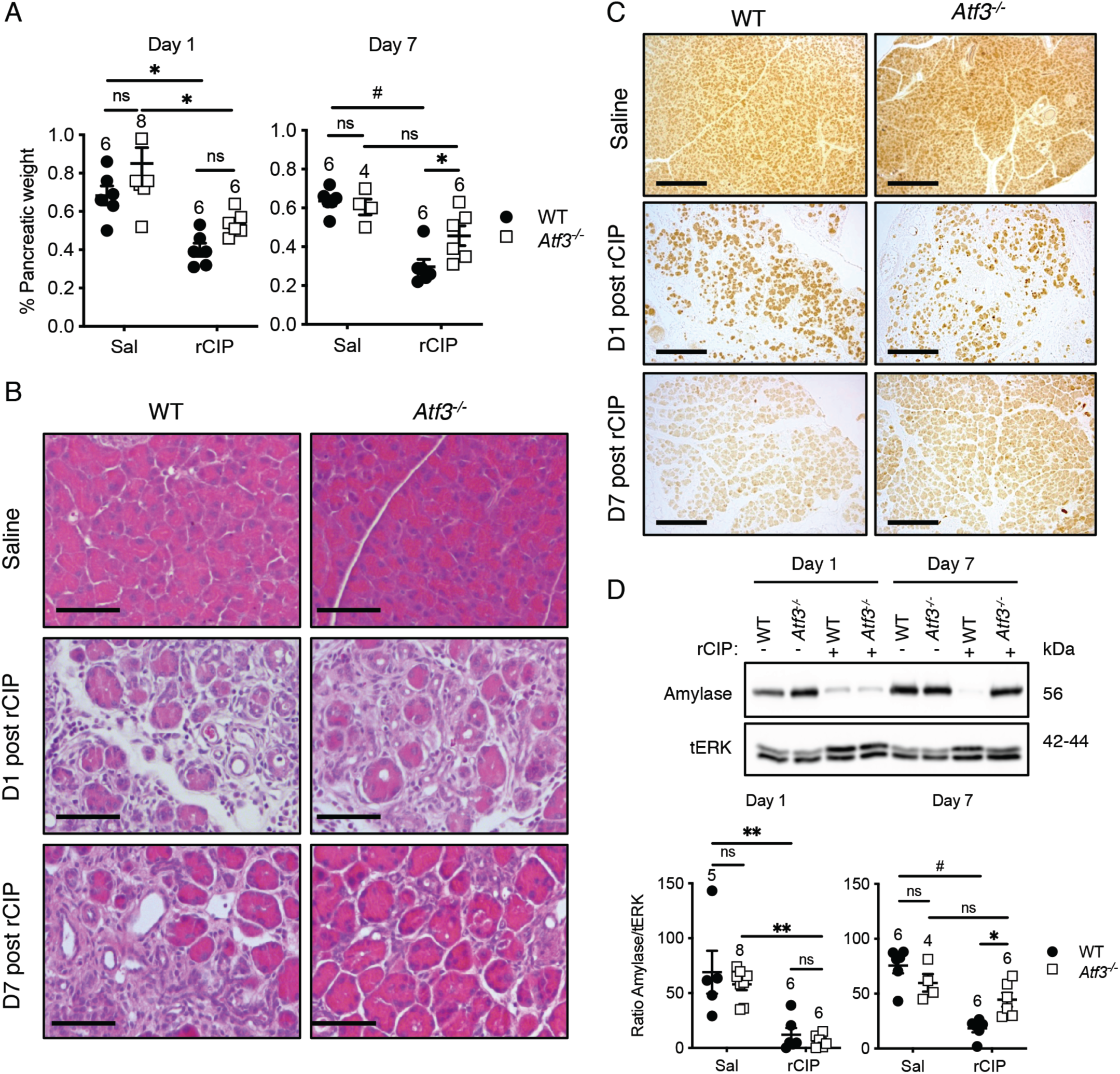
*Atf3*^*-/-*^ mice show accelerated pancreatic regeneration in response to recurrent pancreatic injury. (A) Quantification of pancreatic weight as a % of body weight in mice treated with saline (Sal) or cerulein (rCIP) one and seven days after cessation of treatment. Cerulein-treated wild type (WT) and *Atf3*^*-/-*^ mice showed significant decrease in pancreatic weight at Day 1 with no significant difference between genotypes. Seven days into recovery, only cerulein-treated WT mice showed a decrease in pancreatic weight relative to other groups. (B) Representative H&E histology shows loss of acinar tissue and increased number of duct-like structures one day after rCIP in both genotypes. By day 7 after rCIP, there is reduced damage and increased acinar cell area in *Atf3*^*-/-*^ tissue compared to WT tissue. Magnification bars = 100 μm. (C) Immunohistochemistry (IHC) for amylase in WT and *Atf3*^*-/-*^ pancreatic tissue from mice treated with saline or 1 or 7 days following rCIP. Magnification bar = 400 μm. (D) Western blot analysis and quantification of amylase accumulation in pancreatic extracts from mice treated with saline (-) or rCIP (+). In all cases, ns, not significant; *p<0.05, **p<0.01, #p<0.001; N values are indicated above the data points; error bars represent mean ± SEM. To determine significance, a two-way ANOVA was performed with a Tukey’ post-hoc test.

**Figure 2.**
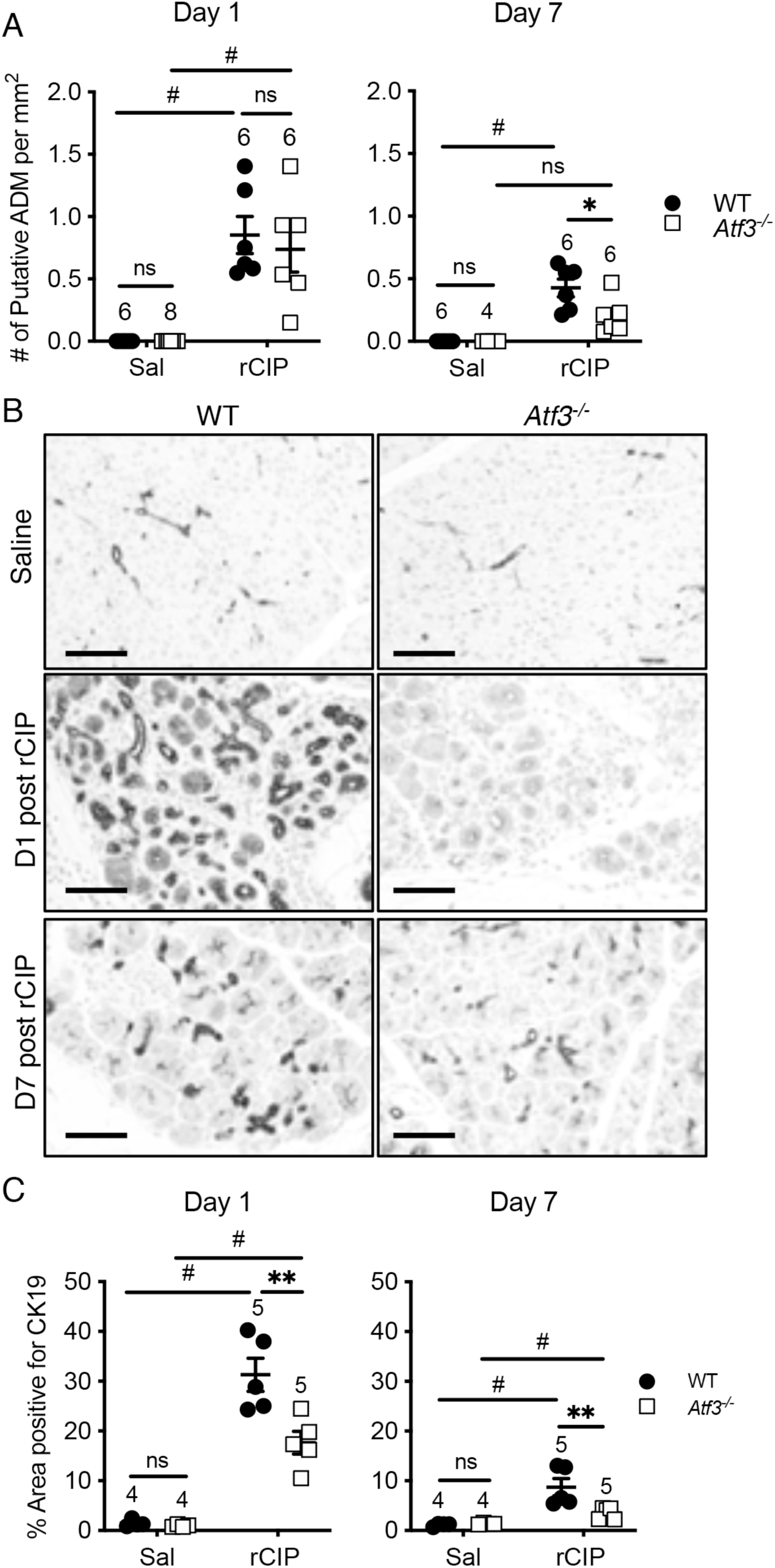
*Atf3*^*-/-*^ mice exhibit reduced ADM in response to recurrent pancreatic injury. (A) Quantification of ADM-like structures following rCIP based on H&E analysis. A significant increase in putative ADM structures was observed one day after rCIP in both genotypes, but no difference between genotypes in the rCIP cohorts. 7 days after rCIP treatment, significantly fewer putative ADM structures were observed in rCIP-treated *Atf3*^*-/-*^ mice compared wild type (WT). (B) Representative IHC for CK19 in WT and *Atf3*^*-/-*^ pancreatic tissue following saline or rCIP. Magnification bars = 70 μm. (C) Quantification of CK19 IHC revealed decreased CK19 accumulation one and seven days after rCIP in *Atf3*^*-/-*^ mice relative to WT mice. In all cases, N values are indicated above the data points; ns, not significant; *p<0.05; **p<0.01, #p<0.001; error bars represent mean ± SEM. To determine significance, a two-way ANOVA was performed with a Tukey’ post-hoc test.

Seven days after rCIP, the differences between WT and *Atf3*^*-/-*^ pancreatic tissue were striking. Pancreas to body weight ratio of cerulein-treated *Atf3*^*-/-*^ mice was similar to saline-treated mice while cerulein-treated WT mice still had significantly smaller pancreata relative to all other groups (Figure 1A). Histological analysis showed almost a complete recovery of acinar tissue in cerulein-treated *Atf3*^*-/-*^ mice while WT mice retained areas of damage (Figure 1B). Acinar tissue was restored more quickly based on histology (Figures 1B, 2A, S2A), amylase accumulation (Figures 1C-E) and CK19 accumulation (Figure 2B, C). Analysis of fibrosis showed no difference between WT and *Atf3*^*-/-*^ mice, with overall fibrosis reduced by day seven compared to day one in both genotypes (Supplemental Figure S2).

Our previous work indicated ATF3 directly regulated expression of transcription factors involved in ADM [22]. IF for SOX9, which is required for ADM [36, 39], showed limited expression in saline-treated animals, localizing specifically to ductal epithelial cells. As previously reported, SOX9 expression was observed in acinar cells and ADM structures 1 and 7 days following rCIP in WT mice (Figure 3A, B). This finding was confirmed by Western blotting (Supplemental Figure S3A). Conversely, SOX9 expression was observed only sporadically in cerulein-treated *Atf3*^*-/-*^ tissue (Figure 3A, B) with no increase observed by Western blot analysis (Supplemental Figure S3A). Similar analysis for PDX1 showed increased accumulation in ADM structures in cerulein-treated WT pancreas tissue, while PDX1 expression was greatly reduced in both intensity (Supplemental Figure S3B) and frequency in cerulein-treated *Atf3*^*-/-*^ pancreas tissue (Supplemental Figure S3C).

**Figure 3.**
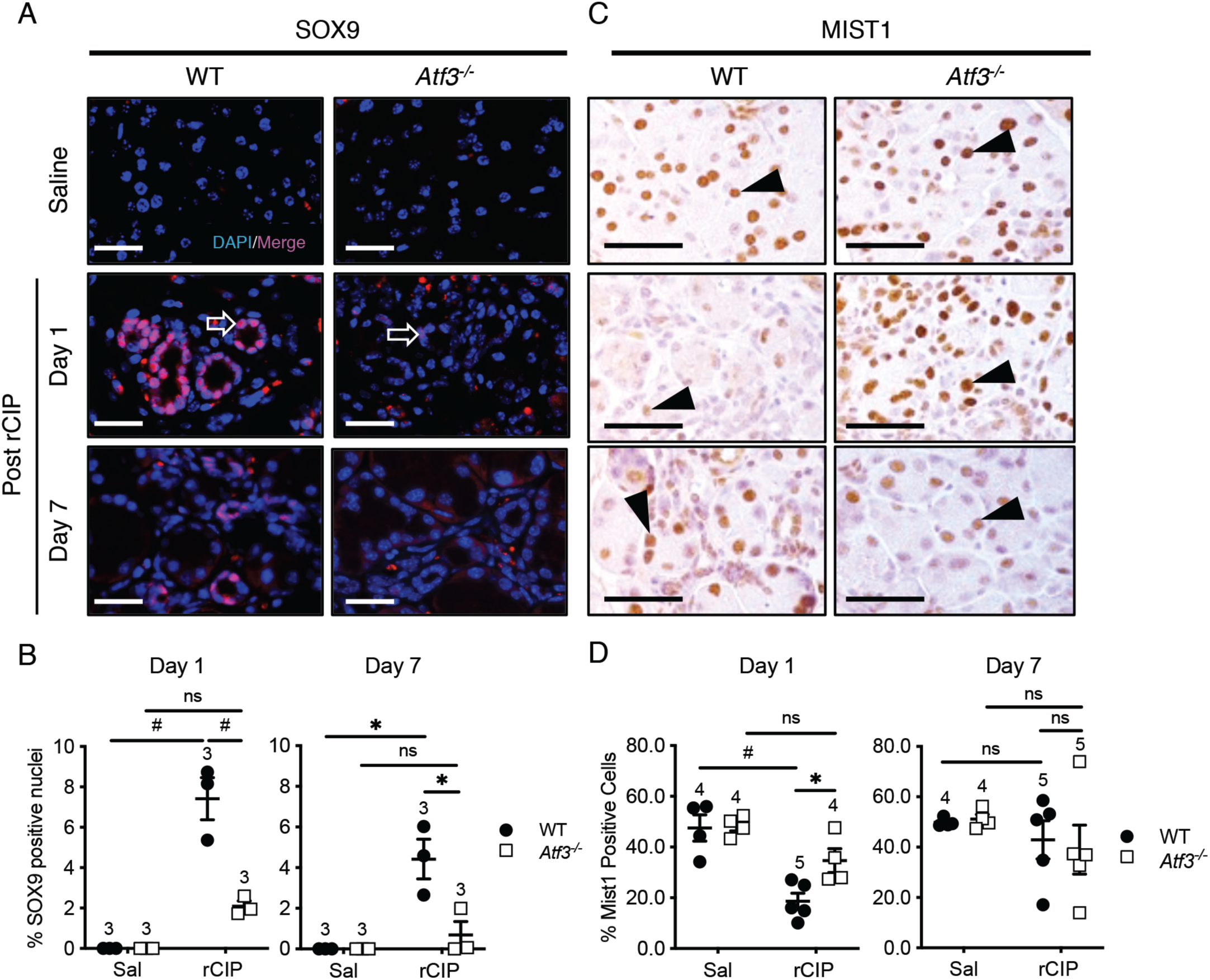
The ADM transcriptional program is reduced in the absence of ATF3. (A) Representative IF for SOX9 in WT and *Atf3*^*-/-*^ pancreatic tissue from mice 1 or 7 days following saline or rCIP treatment. Sections were counterstained with DAPI to reveal nuclei. Arrow indicate SOX9+ cells. Magnification bars = 20 μm. (B) Quantification of the percentage of SOX9+ nuclei showed significantly fewer cells express SOX9 in *Atf3*^*-/-*^ tissue at both time points. (C) Representative IHC for MIST1 in WT and *Atf3*^*-/-*^ pancreatic tissue 1 or 7 days following saline or rCIP. Arrowheads indicate MIST1+ cells. Magnification bars = 50 μm. (D) Quantification of the percent MIST1+ nuclei confirm significantly fewer cells express MIST1 in *Atf3*^*-/-*^ tissue. For graphs, N values are indicated above data points; ns, not significant; *p<0.05, #p<0.001; error bars represent mean ± SEM. To determine significance, a two-way ANOVA was performed with a Tukey’ post-hoc test.

We next examined MIST1, which is required for establishing the mature acinar cell phenotype [40]. As previously reported [22], MIST1 expression is significantly reduced in WT acinar cells one day following cessation of cerulein treatment (Figure 3C, D). Seven days after treatment, MIST1 expression returned to control levels. Conversely, this temporal decrease in MIST1 expression was not observed in cerulein-treated *Atf3*^*-/-*^ tissue (Figure 3C, D). Combined, this data suggests that *Atf3*^*-/-*^ mice recover more quickly from rCIP, possibly through limiting activation of a more progenitor-like state in response to injury.

### Absence of ATF3 reduces spontaneous PanIN progression following KRAS^G12D^ activation

While ATF3 was required for activating the ADM transcriptional program in response to injury, it is possible that oncogenic KRAS can bypass this requirement. Therefore, *Atf3*^*-/-*^ mice were mated to mice allowing inducible activation of oncogenic KRAS (KRAS^G12D^). Recombination was limited to acinar cells by targeting a creERT to the *Ptf1a* gene [41]. These mice contained a *ROSA26* reporter gene (*R26r-YFP*) allowing analysis of cre-mediated recombination. Two to four-month old mice with (*Atf3*^*+/+*^ or *Atf3*^*+/-*^*Ptf1a*^*creERT/+*^*Kras*^*LSL-G12D/+*^; referred to as *Ptf1a*^*creERT/+*^*Kras*^*G12D/+*^) or without ATF3 (*Atf3*^*-/-*^*Ptf1a*^*creERT/+*^*Kras*^*LSL-G12D/+*^; referred to as *APK*) were treated with tamoxifen for five days (Supplemental Figure S4A). This resulted in >97% acinar-specific recombination in both mouse lines with no observable recombination in duct or islet tissue based on activation of the *ROSA26r-LSL-YFP* locus (Supplemental Figure S4B). One cohort of mice was followed for 13 weeks to determine if spontaneous ADM and PanIN lesions were affected by the absence of ATF3 (Supplemental Figure S4A). All groups showed a decrease in weight in response to tamoxifen, which was regained over the course of the experiment (Supplemental Figure S4C, D). Upon dissection, 2/5 *Ptf1a*^*creERT/+*^*Kras*^*G12D/+*^ mice contained pancreata with some fibrotic masses observed at the gross morphological level (data not shown). Pancreatic weight as a percentage of total body weight was increased in *Ptf1a*^*creERT/+*^*Kras*^*G12D/+*^ mice relative to all other genotypes (Figure 4A). H&E analysis confirmed significant disruptions to pancreatic architecture, including loss of acini, extensive PanIN lesions, inflammation, and fibrosis in 2/5 *Ptf1a*^*creERT/+*^*Kras*^*G12D/+*^ mice (Figure 4B). In the remaining three *Ptf1a*^*creERT/+*^*Kras*^*G12D/+*^ mice, areas of ADM and early PanIN lesions were readily observed (Figure 4C). IF revealed SOX9 expression within the ADM and PanIN lesions observed in *Ptf1a*^*creERT/+*^*Kras*^*G12D/+*^ tissue (Figure 4D). Conversely, *APK* mice showed only sporadic ADM/PanIN lesions (Figure 4B) with minimal SOX9 expression (Figure 4D) and no PanIN2 or PanIN3 lesions. These results indicated ATF3 was required for spontaneous KRAS^G12D^-mediated PanIN formation.

**Figure 4.**
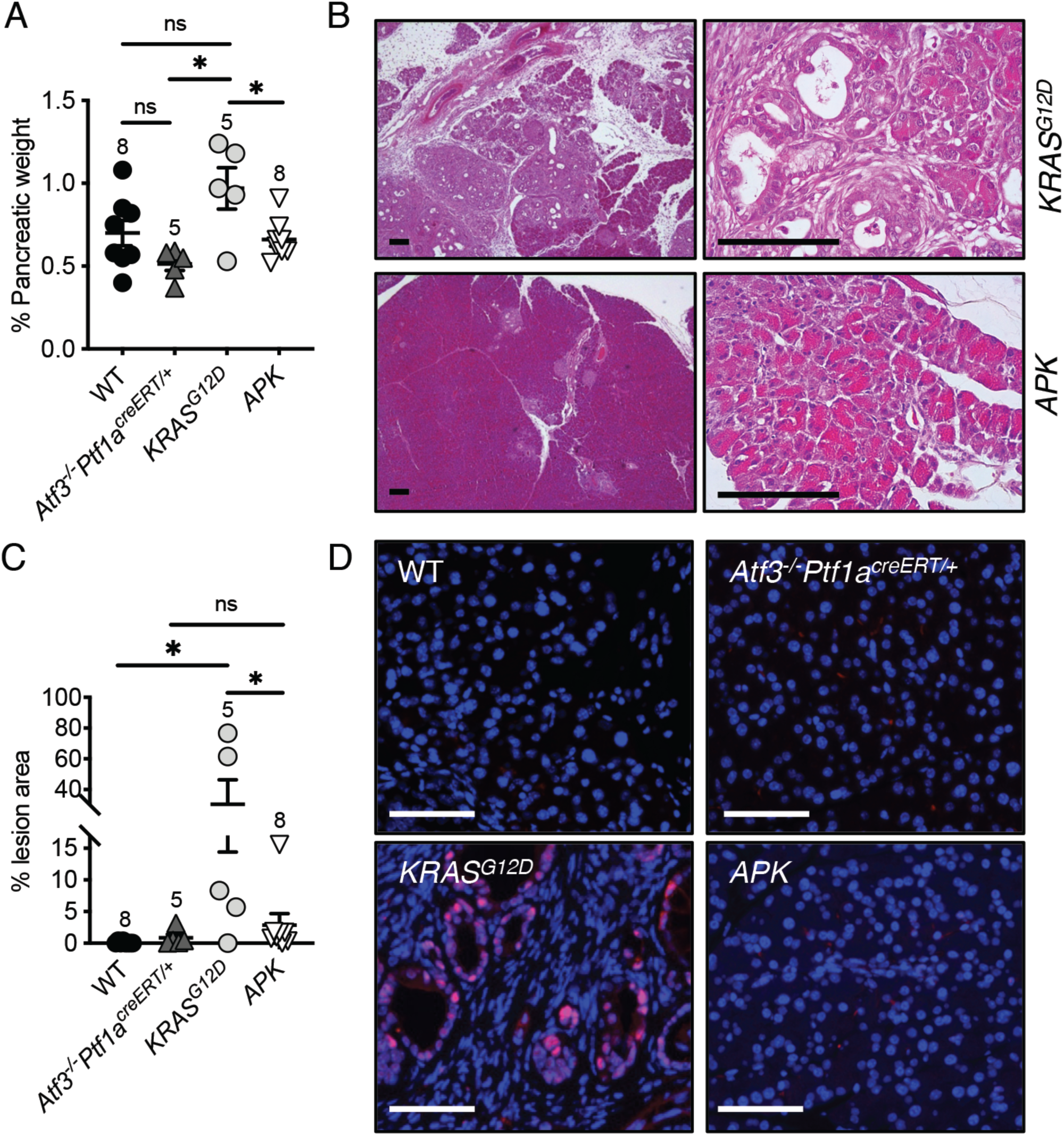
*Atf3*^*-/-*^ mice show minimal spontaneous ADM and PanIN formation following activation of oncogenic KRAS. (A) Quantification of pancreatic weight as a % of body weight 13 weeks after activation of KRAS^G12D^ with tamoxifen. Pancreatic weight is increased in *Ptf1a*^*creERT/+*^*Kras*^*G12D/+*^ (indicated as KRAS^G12D^) mice relative to wild type (WT), *Ptf1a*^*creERT/+*^*Atf3*^*-/-*^ and *APK* mice. (B) Representative H&E on pancreatic tissue from *Ptf1a*^*creERT/+*^*Kras*^*G12D/+*^ and *APK* mice 13 weeks after KRAS^G12D^ activation. Extensive lesions were observed in 2/5 *Ptf1a*^*creERT/+*^*Kras*^*G12D/+*^ mice but not in any other genotype. Magnification bars = 100 μm. (C) Quantification of lesion area based on H&E histology. Lesion area was increased in *Ptf1a*^*creERT/+*^*Kras*^*G12D/+*^ tissue relative to all other genotypes. For all graphs, N values are indicated above data points; ns, not significant; *p<0.05; error bars represent mean ± SEM. To determine significance, a one-way ANOVA was performed with a Tukey’ post-hoc test. (D) Representative IF for SOX9 (red) on pancreatic tissue for each genotype. Sections were counterstained with DAPI to reveal nuclei. Magnification bars = 60 μm.

### Oncogenic KRAS bypasses the requirement of ATF3 for ADM following injury

Next, we exposed *Ptf1a*^*creERT/+*^*Kras*^*G12D/+*^ and *APK* mice to a two-day regime of acute cerulein treatment seven days after tamoxifen treatment and examined pancreatic tissue two and five weeks after cerulein treatment (Supplemental Figure S4A). During tamoxifen treatment and acute CIP, mice were weighed daily. All saline-treated mice, regardless of ATF3 expression or oncogenic KRAS activity gained body weight over the experimental time period (Supplemental Figure S5A). CIP-treated *Ptf1a*^*creERT/+*^*Kras*^*G12D/+*^ mice showed reduced body weight starting 43 days after tamoxifen treatment compared to *Ptf1a*^*creERT/+*^ and *Atf3*^*-/-*^*Ptf1a*^*CreERT/+*^ mice, while *APK* mice treated with cerulein had significantly reduced body weight compared to all other genotypes 31 days into treatment (Supplemental Figure S5B). Both *Ptf1a*^*creERT/+*^*Kras*^*G12D/+*^ (3/22) and *APK* lines (1/14) exhibited some mortality during the 5-week experimental time point. Upon dissection, *APK* mice appeared much smaller than other groups with negligible abdominal wall muscle observed in most of these mice (data not shown). *APK* mice also had significantly smaller pancreata at both time points examined relative to other genotypes (Supplementary Figure S5C, D). Conversely, cerulein-treated *Ptf1a*^*creERT/+*^*Kras*^*G12D/+*^ mice had significantly increased pancreatic weight relative to body weight (Supplementary Figure S5D) compared to all other groups two weeks post treatment, with multiple fibrotic nodules visible (Supplementary Figure S5C). By five weeks, the difference in pancreatic/body weight was no longer apparent in *Ptf1a*^*creERT/+*^*Kras*^*G12D/+*^ mice, even though nodules were still present. Histological analysis of pancreatic tissue indicated widespread loss of acinar tissue and development of PanIN lesions within two weeks of inducing CIP in *APK* and *Ptf1a*^*creERT/+*^*Kras*^*G12D/+*^ pancreatic tissues (Figure 5A). The loss of acinar tissue was confirmed by a complete absence of amylase accumulation based on western blot analysis (Figure 6A, B).

**Figure 5.**
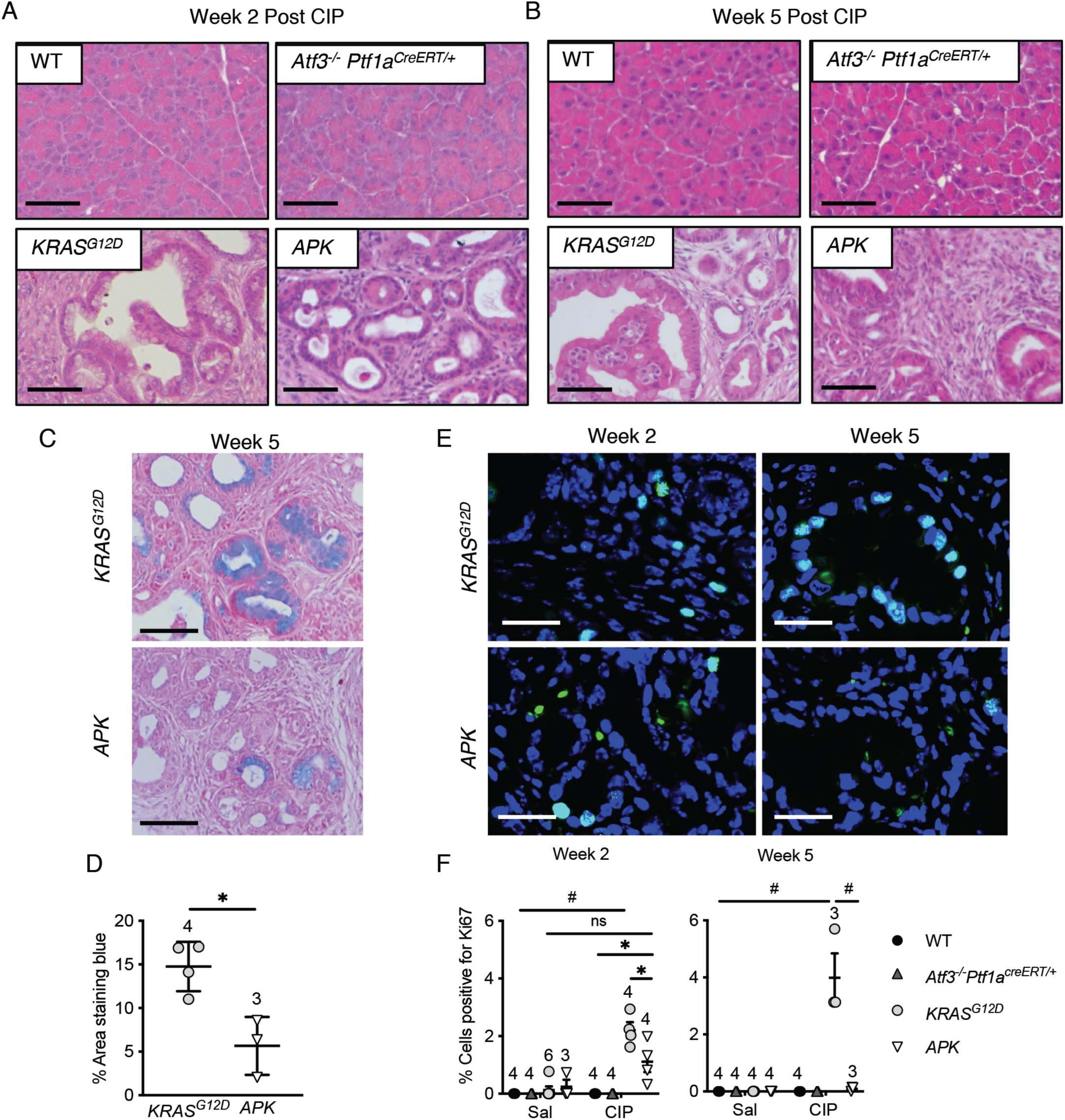
ATF3 is dispensable for progression but not maintaining PanINs when KRAS^G12D^ activation is coupled with pancreatic injury. Representative H&E histology on pancreatic tissue (A) two and (B) five weeks after cerulein treatment for all genotypes. No lesions were observed in WT or *Atf3*^*-/-*^*Ptf1a*^*creERT/+*^ tissue. Extensive lesions and no acinar tissue was visible at either time point in both *Ptf1a*^*creERT/+*^*Kras*^*G12D/+*^ (KRAS^G12D^) and *APK* tissue following cerulein treatment. Magnification bars = 100 μm. (C) Representative Alcian blue histology on *Ptf1a*^*creERT/+*^*Kras*^*G12D/+*^ and *APK* tissue following cerulein treatment showed significantly more alcian blue staining in *Ptf1a*^*creERT/+*^*Kras*^*G12D/+*^ mice which is quantified in (D). (E) Representative IF for Ki-67 in *Ptf1a*^*creERT/+*^*Kras*^*G12D/+*^ and *APK* tissue 2 and 5 weeks following cerulein treatment. Magnification bar = 50 μm. (F) Quantification of the percentage of Ki-67+ cells indicated significantly fewer positive cells in *APK* tissue. For all graphs, N values are indicated in brackets or above data points; ns, not significant; *p<0.05, #p<0.001; error bars represent mean ± SEM. To determine significance, a student’s t-test (D) or (F) two-way ANOVA was performed with a Tukey’ post-hoc test.

**Figure 6.**
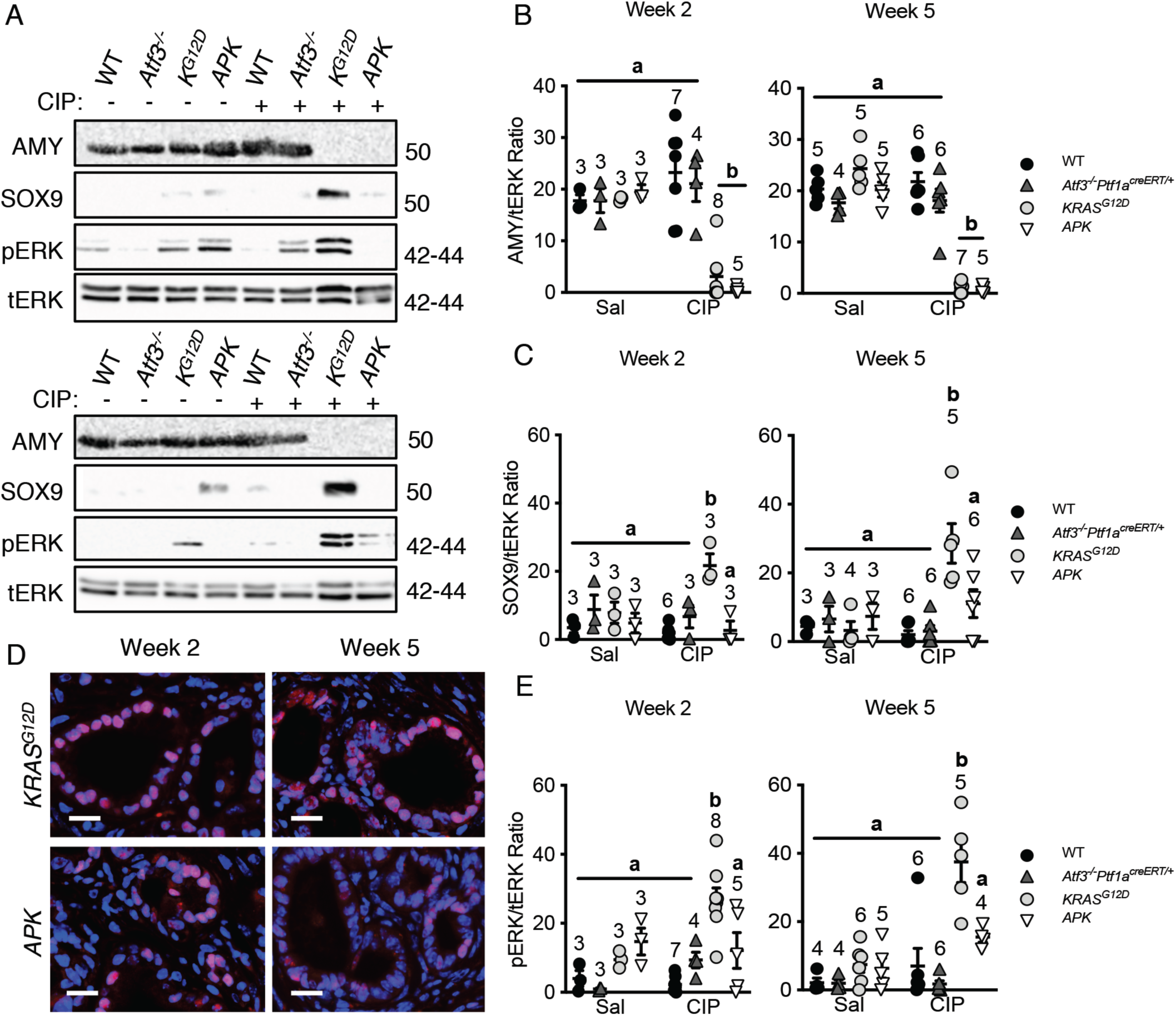
ATF3 is required for complete establishment and maintenance of the molecular ADM profile in the presence of KRAS^G12D^. (A) Representative western blots for amylase (AMY), SOX9, phosphorylated ERK (pERK) and total ERK (tERK, loading control). No detectable amylase accumulation is observed in *Ptf1a*^*creERT/+*^*Kras*^*G12D/+*^ or *APK* tissue (quantified in B) while SOX9 and pERK accumulation increases only in *Ptf1a*^*creERT/+*^*Kras*^*G12D/+*^ tissue (quantified in C and E, respectively). Similar results are obtained both 2 and 5 weeks after cerulein (CIP) treatment. For all graphs, N values are indicated above data points; significantly different values are indicated by letters. In (C) and (E), *Ptf1a*^*creERT/+*^*Kras*^*G12D/+*^ values are significantly different from all genotypes (p<0.001) even from *APK* mice (p<0.05). Error bars represent mean ± SEM. To determine significance, a two-way ANOVA was performed with a Tukey’ post-hoc test. (D) Representative IF for SOX9 (red) on pancreatic tissue from *Ptf1a*^*creERT/+*^*Kras*^*G12D/+*^ or *APK* mice. Sections were counterstained with DAPI to reveal nuclei. Magnification bars = 20 μm.

Cursory H&E analysis suggests ATF3 is not required for KRAS^G12D^-mediated PDAC progression. Saline and cerulein-treated control *Ptf1a*^*creERT/+*^ and *Atf3*^*-/-*^*Ptf1a*^*creERT/+*^ tissue showed no damage either two or five weeks after treatment (Supplemental Figure 6A; Figure 5A). To determine if the pathological phenotype was the same between *APK* and *Ptf1a*^*creERT/+*^*Kras*^*G12D/+*^ mice, tissue was assessed based on the highest grade lesion present per tissue area (Table 1). As suggested from 13-week analysis, saline-treated *APK* and *Ptf1a*^*creERT/+*^*Kras*^*G12D/+*^ mice showed some pre-neoplastic lesions (Supplemental Figure S6A, B; Table 1). Upon CIP-treatment, *APK* and *Ptf1a*^*creERT/+*^*Kras*^*G12D/+*^ mice treated with cerulein showed high grade PanIN3 lesions two weeks after CIP (Figure 5A; Table 1). However, by week five, *APK* tissue showed predominately low grade PanIN1 and 2 lesions compared to predominantly high grade PanIN3 and PDAC lesions observed in *Ptf1a*^*creERT/+*^*Kras*^*G12D/+*^ tissue (Table 1). The decrease in progressive PanIN lesions in *APK* mice was also reflected by significantly lower accumulation of CK19+ ducts in *APK* tissue compared to *Ptf1a*^*creERT/+*^*Kras*^*G12D/+*^ mice (Supplementary Figure S6B, C). Alcian blue stain, which identifies mucin and suggests metaplastic PanIN lesions, was significantly reduced in *APK* tissue relative to *Ptf1a*^*creERT/+*^*Kras*^*G12D/+*^ tissue (Figure 5C, D). Finally, epithelial cell proliferation in PanINs was examined by Ki-67 fluorescence. Ki67+ cells were reduced at two weeks and completely absent at five weeks in *APK* lesions relative to *Ptf1a*^*creERT/+*^*Kras*^*G12D/+*^ tissue (Figure 5E, F). Therefore, while overall histology was similar between *APK* and *Ptf1a*^*creERT/+*^*Kras*^*G12D/+*^ tissue, multiple analyses suggest the absence of ATF3 affects KRAS^G12D^-mediated progression to PDAC.

**Table 1.** Histological analysis of pancreatic lesions in response to saline or cerulein treatment^1^

|  | *Ptf1a<sup>creERT/+</sup>* |  | *Atf3<sup>-/-</sup>Ptf1a<sup>creERT/+</sup>* |  | *Ptf1a<sup>creERT/+</sup>KRAS<sup>G12D/+</sup>* |  | *APK* |  |
| --- | --- | --- | --- | --- | --- | --- | --- | --- |
| Saline | 2 weeks | 5 weeks | 2 weeks | 5 weeks | 2 weeks | 5 weeks | 2 weeks | 5 weeks |
| Normal | 4 | 2 | 3 | 2 | 1 | 0 | 1 | 1 |
| ADM | 0 | 1 | 0 | 0 | 0 | 0 | 1 | 0 |
| PanIN1 | 0 | 0 | 0 | 0 | 2 | 2 | 1 | 1 |
| PanIN2 | 0 | 0 | 0 | 0 | 0 | 1 | 0 | 1 |
| PanIN3 | 0 | 0 | 0 | 0 | 0 | 0 | 0 | 0 |
| PDAC | 0 | 0 | 0 | 0 | 0 | 0 | 0 | 0 |
| Total # of mice | 4 | 3 | 3 | 2 | 3 | 3 | 3 | 3 |
| Cerulein |  |  |  |  |  |  |  |  |
| Normal | 0 | 2 | 2 | 0 | 0 | 0 | 0 | 0 |
| ADM | 2 | 0 | 0 | 1 | 0 | 0 | 0 | 0 |
| PanIN1 | 0 | 0 | 1 | 0 | 0 | 0 | 2 | 1 |
| PanIN2 | 0 | 0 | 0 | 0 | 2 | 2 | 0 | 5 |
| PanIN3 | 0 | 0 | 0 | 0 | 9 | 4 | 4 | 2 |
| PDAC | 0 | 0 | 0 | 0 | 1 | 1 | 0 | 0 |
| Total # of mice | 2 | 2 | 3 | 1 | 12 | 7 | 6 | 8 |

<sup>1</sup> – pancreatic tissue was scored for the highest grade lesion within the tissue. Numbers do not include three
*Ptf1a<sup>creERT/+</sup>KRAS<sup>G12D/+</sup>* and one *APK* mouse that needed to be sacrificed prior to the experimental end point

We next examined if the molecular profile for ADM progression was present in *APK* mice. Both transcriptional (SOX9) and signaling mediators (phosphorylated MAPK1/2) of ADM (**Figure 6**) were examined. IF for SOX9 showed equivalent numbers of SOX9+ cells two weeks after CIP in *Ptf1a*^*creERT/+*^*Kras*^*G12D/+*^ and *APK* tissue (Figure 6D), but with significantly lower SOX9 accumulation in *APK* tissue based on western blot analysis (Figure 6A, C; SOX9 accumulation is 7.99 ± 1.24-fold higher in *Ptf1a*^*creERT/+*^*Kras*^*LSL-G12D/+*^ extracts compared to *APK*). By five weeks post-cerulein, both the number (Figure 6D) and level of SOX9 accumulation was reduced in *APK* mice (Figure 6C; 2.58 ± 1.41-fold higher in *Ptf1a*^*creERT/+*^*Kras*^*LSL-G12D/+*^ extracts). The decreased expression of the ADM-promoting SOX9 was consistent with decreased activation of MAPK signalling in *APK* tissue based on western blot analysis for pERK both two (2.21 ± 0.66-fold less pERK) and five weeks (2.42 ± 0.25-fold less pERK) after injury compared to *Ptf1a*^*creERT/+*^*Kras*^*G12D/+*^ tissue (Figure 6A, E). Combined, these results suggest that the absence of ATF3 in the presence of KRAS^G12D^ reduces the activation of factors promoting ADM and PanIN formation. Therefore, while the requirement of ATF3 for ADM can be bypassed by KRAS^G12D^, ATF3 appears to be required for maintaining high-grade PanIN lesions.

### An absence of ATF3 affects the fibrotic and inflammatory pathways affected by oncogenic KRAS

Previous analysis of ATF3 in other cancers suggests an important role in fibrosis and inflammatory processes. To determine if the global absence of ATF3 affects these two elements in PDAC progression, we compared expression of markers for stellate cells and inflammation between *APK* and *Ptf1a*^*creERT/+*^*Kras*^*G12D/+*^ mice. Fibrosis, as determined by Mason’s Trichrome stain, was significantly higher two weeks post injury in *APK* tissue compared to *Ptf1a*^*creERT/+*^*Kras*^*G12D/+*^ tissue (Figure 7), suggesting accelerated fibrosis in the absence of ATF3. By five weeks, no difference was observed in fibrosis between these two genotypes (Figure 7) as the level of fibrosis appears to lessen in *APK* mice. Interestingly, analysis of α-smooth muscle actin (α-SMA), a marker of activated fibroblast cells, showed no difference in accumulation based on western blot analysis (Supplementary Figure 7).

**Figure 7.**
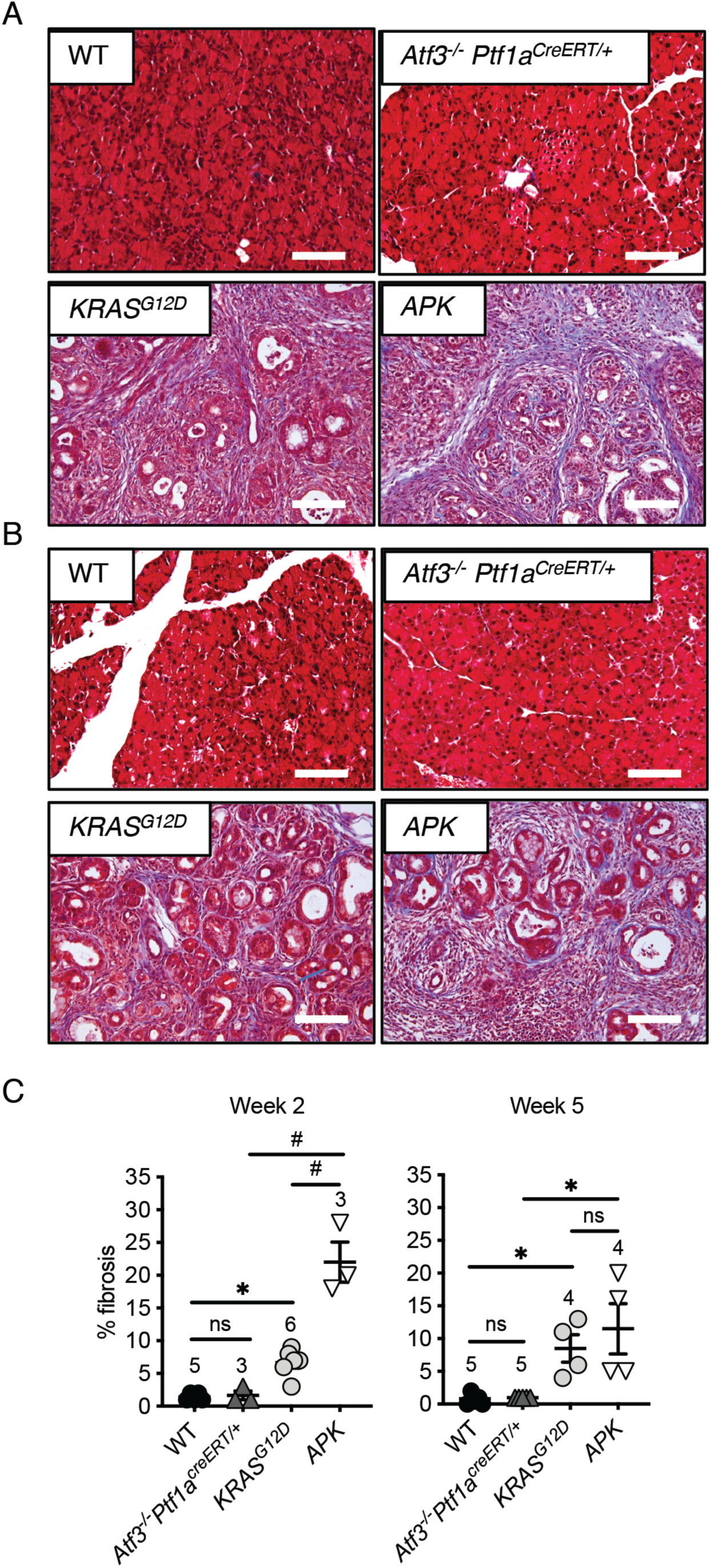
*APK* mice display enhanced fibrosis at early time point after pancreatic injury. Representative images of Mason trichrome staining (A) two or (B) five weeks after cerulein treatment in all genotypes. Magnification bars = 140 µm. (C) Quantification of fibrosis based on trichrome staining, showed increased fibrosis in *APK* tissue compared to all other genotypes two weeks post-CIP and increased fibrosis in *Ptf1a*^*creERT/+*^*Kras*^*G12D/+*^ (*KRAS*^*G12D*^) and *APK* mice five weeks after cerulein treatment. N values are indicated above data points; ns, not significant; *p<0.05, #p<0.001; error bars represent mean ± SEM. To determine significance, a one-way ANOVA was performed with a Tukey’ post-hoc test.

Next, to compare the immune cell infiltrate between *APK* and *Ptf1a*^*creERT/+*^*Kras*^*G12D/+*^ mice, we performed mass cytometry (CyTOF) on pancreatic tissue from mice two weeks after cerulein treatment. CyTOF experiments revealed a decreased infiltration of macrophages in *APK* mice with a concomitant increase in T lymphocyte accumulation (Figure 8A, B). To confirm these findings, we performed IF for F4/80, a macrophage-specific marker (Figure 8C). In confirmation of the CyTOF experiments, significantly fewer macrophages were observed in *APK* tissue compared to *Ptf1a*^*creERT/+*^*Kras*^*LSL-G12D/+*^ tissue five weeks after injury (Figure 8D). These results suggest the global loss of ATF3 has both acinar and non-acinar specific role in affecting KRAS^G12D^-mediated PDAC progression.

**Figure 8.**
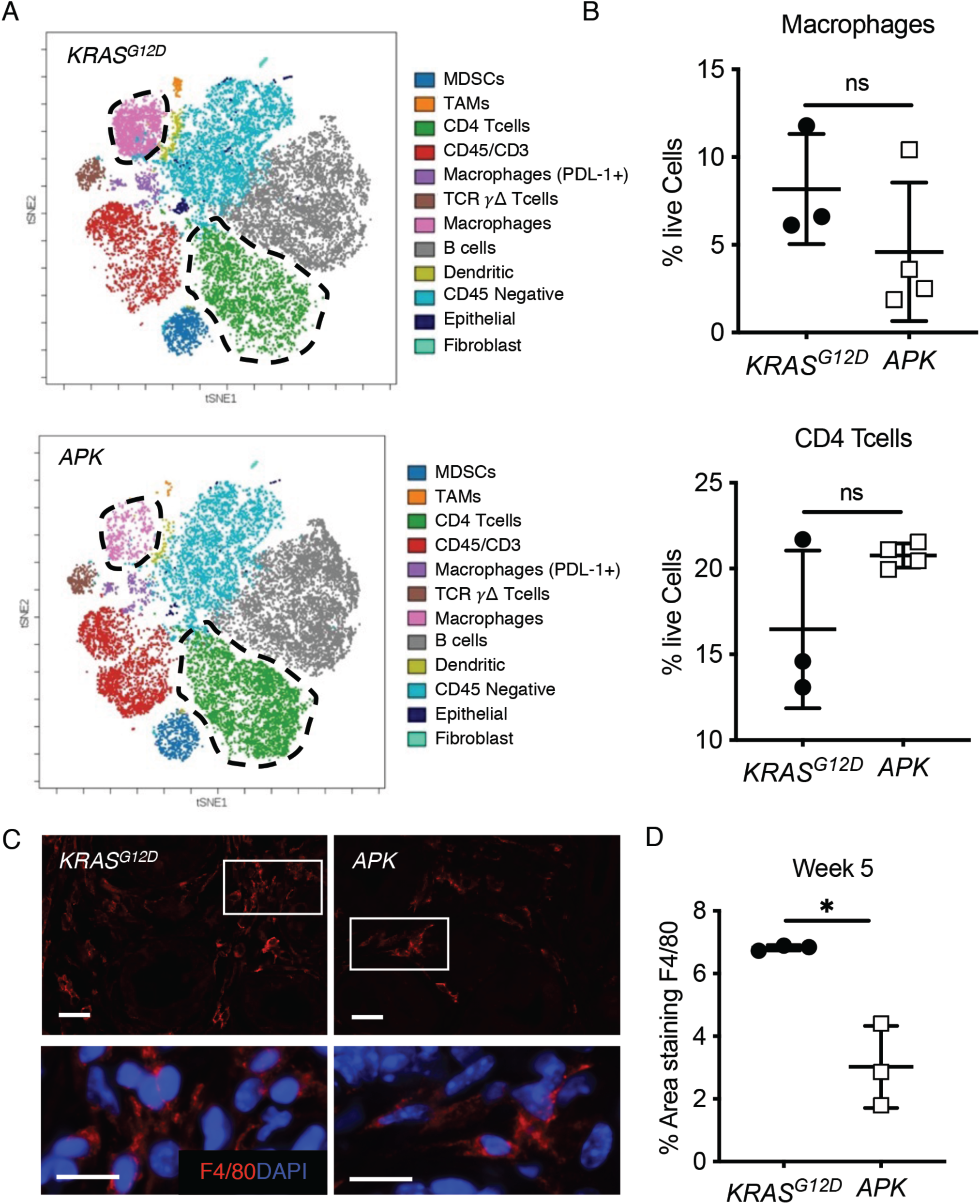
APK mice showed an altered inflammatory response to pancreatic damage. (A) Representative viSNE analysis performed on cytometric flow data obtained from *Ptf1a*^*creERT/+*^*Kras*^*G12D/+*^ (*KRAS*^*G12D*^) and *APK* mice pancreata at 2 weeks post CIP (n=3-4). (B) Quantification of macrophage and CD4 T cells based on viSNE analysis suggested decreased macrophage and increased T lymphocyte accumulation in *APK* pancreata relative to *Ptf1a*^*creERT/+*^*Kras*^*G12D/+*^. (C) Representative IF images show significant fewer macrophages based on the percent area or F4/80 staining F4/80 in *APK* mice, which is quantified in (D). *P<0.05, error bars represent % mean ± SEM; N=3. To determine significance, a student’s t-test was performed.

## Discussion

In this study, we identified a novel role for ATF3 in recurrent pancreatic injury and progression to PDAC. Recurrent pancreatitis is a significant risk factor for PDAC, possibly due to the combined effects of inflammation and loss of the acinar cell phenotype through ADM. Through examining the effects of *Atf3* deletion on the acinar cell response to recurrent injury or constitutive KRAS activation, we identified a complex role for ATF3 in affecting multiple cell types within the pancreas. The absence of ATF3 prevented full activation of the ADM transcriptional profile and limited spontaneous PanIN formation in the presence of oncogenic KRAS. This requirement for ATF3 in ADM and PanIN progression appears dispensable when constitutive KRAS activation is combined with injury. However, the maintenance of high grade PanINs and progression to PDAC still required ATF3. In addition, we observed a transient increase in fibrosis and an altered inflammatory response with a decreased macrophage infiltration. These results suggest that, in addition to ADM, ATF3 also affects stellate cells and myeloid cells. Whether this is through cell autonomous pathways or mediated by epithelial cell interactions needs to be investigated in future studies.

Previous work from our lab identified a requirement for ATF3 in activating ADM transcriptional profile during acute injury. We have now assessed ATF3’s role in conditions that promote PDAC. As observed in acute injury, ATF3 is required for changing the expression of key transcriptional regulators of the ADM process, including PDX1, SOX9 and MIST1. Increased expression of SOX9 and PDX1 observed during injury [42] is required for maintaining ADM morphology and expression of duct-specific genes such as *Ck19* [43]. Conversely, MIST1 expression is rapidly decreased. In the absence of ATF3, SOX9 and PDX1 expression are reduced compared to WT mice, while MIST1 expression was not completely repressed. The incomplete activation of the ADM transcriptome likely accounted for the more rapid regeneration exhibited by *Atf3*^*-/-*^ mice following rCIP. Interestingly, structures resembling ADM were still observed and amylase accumulation was almost completely lost even while some MIST1 expression persisted. Therefore, recurrent injury still promoted the loss of mature acinar cell phenotype in the absence of ATF3, even without activation of SOX9. Similar observations of partial ADM have been observed in *Sox9*-deficient acinar cells [44], likely through compensatory mechanisms involving Hepatocyte nuclear factor 6 (HNF6; [45]. HNF6 works synergistically with SOX9 to promote ADM development.

Based on the rapid regeneration and reduced ADM exhibited by *Atf3*^*-/-*^ mice, as well as the finding that *APK* mice do not develop PDAC, it is tempting to suggest ATF3 may be a target for inhibiting KRAS-mediated PDAC. In fact, we observed little to no transformation of acinar cells following activation of KRAS^G12D^ in mature *APK* acinar cells. Conversely, high grade PanINs were readily observed in a subset of *Ptf1a*^*creERT/+*^*Kras*^*LSL-G12D/+*^ mice 13 weeks after KRAS^G12D^ activation. However, our findings suggest the absence of ATF3 is detrimental to regeneration when oncogenic KRAS is expressed in addition to injury. Following induction of pancreatitis, acinar cells expressing KRAS^G12D^ appear to bypass the requirement of ATF3. SOX9 expression increases and high grade PanIN lesions are observed in *APK* mice two weeks following injury. However, SOX9 accumulation was reduced in *APK* mice two weeks following injury and almost completely absent by five weeks suggesting an altered molecular profile for ADM and PanINs in the absence of ATF3. Unlike SOX9, the expression of *Hnf6* is not maintained in more progressive lesions (PanIN lesions; [45], which may explain the inability of *APK* tissue to progress to PDAC. These findings support histological findings that indicate a reduction in high-grade lesions in *APK* tissue.

While these findings support ATF3 requirement for persistent ADM, unlike in rCIP, the pancreata in *APK* mice did not regenerate. *APK* pancreata were significantly smaller in size than all other genotypes and histological analysis revealed increased fibrosis in *APK* tissue at two weeks, suggesting *Atf3*’s deletion affects stellate cell function. Whether the fibrosis observed in *APK* tissue limits pancreatic regeneration in these mice is unclear, but the amount and type of fibrosis can affect progression to PDAC. There is also a loss of fibrosis between two and five weeks in *APK* mice. While it is likely that this results from a general wasting within *APK* mice, we cannot rule out the possibility that factors are promoting resorption of the ECM. Therefore, future work will assess the ECM to determine if stellate cells in *APK* mice have a different expression profile. We also observed a cachexia-like phenotype in *APK* mice, which lost on average 15% of their starting body weight within five weeks. Cachexia results in part from an altered metabolic phenotype, leading to significant loss of muscle. While ATF3 has not yet been implicated in metabolism, the related protein ATF4 does regulate amino acid metabolism in CD4+ T cells. Our previous work showed ATF3 targeted many genes also regulated by ATF4 and Gene Ontology analysis identified dysregulation of several pathways involved in amino acid metabolism in *Atf3*^*-/-*^ mice in response to acute pancreatic injury [22]. In support of a role for ATF3 in the inflammatory response during PDAC, we observe decreased myeloid cell infiltration combined with a trend in increased CD4+ T cells. Whether the gene expression/function of these cells is altered needs to be assessed.

Ultimately, this study was limited by the use of a germ line deletion of *Atf3*. We chose to use these mice as they show no phenotype without the induction of some form of stress. However, we cannot determine whether the phenotypes are the result of acinar or non-acinar requirements for ATF3 gene regulation. We previously showed ATF3 enrichment at ∼35% of the genes altered four hours into cerulein-induced pancreatic injury including genes involved in affecting metabolism, promoting inflammation and ECM production. Therefore, it is possible that ATF3 regulates stellate and inflammatory cells indirectly through cross-talk with acinar cells. Lineage tracing analysis confirmed recombination, and therefore KRAS^G12D^ expression, was limited to acinar cells and PanIN lesions derived from acinar cells. Therefore, we are not observing the results of activating KRAS in non-acinar cells. It is known that ATF3 can affect cancer progression in other systems through non-cell autonomous regulation [46-48]. However, these studies identified a non-cell autonomous for non-tumour cells expressing ATF3. Published data identified ATF3 expression in both inflammatory and ECM-producing cells in other pathologies including breast and lung cancer [29, 49] and our findings indicate ATF3 is expressed in these cell populations in the context of PDAC. Therefore, it is likely that ATF3 directly affects the function of both stellate and myeloid cells in the context of PDAC. Indeed, it is possible that the transient increase in fibrosis and inability to maintain high grade lesions in *APK* mice may be due to the difference in inflammatory response. Tumor-associated macrophages promote cancer fibrosis by secreting factors that activate fibroblast-mediated extracellular matrix remodeling [50].

In summary, the findings in this study support several roles for ATF3 in pancreatic injury and PDAC related to acinar, stellate and immune cells. Future work to tease out cell-specific roles for ATF3 will need to involve cell-specific deletion of *Atf3* in the context of oncogenic KRAS. In addition, the potential for targeting the UPR, and ATF3 specifically, in the context of pancreatic pathologies will need to account for this multifaceted role.

## Materials and Methods

### Mouse Models

Two-to-four month old male and female C57/Bl6 mice or congenic mice carrying a germline deletion of *Atf3* (*Atf3*^*-/-*^; [51]) were used for recurrent cerulein-induced pancreatitis (rCIP) studies. Alternatively, *Atf3*^*-/-*^ mice were bred to mice in which a tamoxifen-inducible *cre* recombinase (*creERT*) was targeted to the *Ptf1a* allele (*Ptf1a*^*creERT/+*^; [41]. *Atf3*^*-/-*^*Ptf1a*^*creERT/+*^ mice were crossed to mice carrying a constitutively active *Kras* gene (KRAS^G12D^) preceded by *loxP* sites flanking a stop codon (*loxP-stop-loxP*; *LSL*) and targeted to the *Kras* allele (*Kras*^*LSL-G12D/+*^; [52]. Through subsequent mating on a C57/Bl6 background, we generated *Atf3*^*+/+*^*Ptf1a*^*creERT/+*^*Kras*^*LSL-G12D/+*^ (referred to as *Ptf1a*^*creERT/+*^*Kras*^*LSL-G12D/+*^) and *Atf3*^*-/-*^ *Ptf1a*^*creERT/+*^*Kras*^*LSL-G12D/+*^ mice (referred to as *APK*). In some cases, mice heterozygous for *Atf3* (*Atf3*^*+/-*^) were included in the *Ptf1a*^*creERT/+*^*Kras*^*LSL-G12D/+*^ cohort as loss of a single copy of *Atf3* has no documented effects REF. To allow lineage tracing of acinar cells, *Ptf1a*^*creERT/+*^*Kras*^*LSL-G12D/+*^ and *APK* mice were mated to mice containing a *yellow fluorescent protein* (*YFP*) gene downstream of a *LSL* cassette targeted to the *Rosa26r* (*Rosa26r*^*LSL-YFP/+*^) allele. Genotypes were confirmed before and after experimentation using the primers indicated in Supplementary Table S1.

### Cerulein-induced pancreatitis

Mice were given normal chow and water *ad libitum* throughout the experiment. To induce recurrent pancreatic injury, *Atf3*^*-/-*^ and *Atf3*^*+/+*^ mice received intraperitoneal injections of cerulein (250 µg/kg body weight; Sigma; Cat. #17650-98-5; St. Louis, MO) or 0.9% saline (control) twice daily (9 00 h and 15 00 h) for 14 days (Supplemental Figure S1). Mice were weighed daily to determine changes in body weight. Mice were killed one or seven days after the last cerulein injections. Pancreatic weight (g) was determined post mortem and compared to total body weight.

For experiments involving KRAS^G12D^, *Atf3*^*+/+*^*Ptf1a*^*creERT/+*^ (designated wild type; WT); *Atf3*^*-/-*^*Ptf1a*^*creERT/+*^, *Ptf1a*^*creERT/+*^*Kras*^*LSL-G12D/+*^ and *APK* mice received 5 mg of tamoxifen (Sigma-Aldrich; Cat. #10540-29-1) each day for 5 days via oral gavage, producing cre-recombination efficiency >95% [41]. Seven days following tamoxifen treatment, cerulein (50 µg/kg) was administered via IP injection; 8 times over 7 hours (n values are indicated in each figure). Mice were weighed daily to monitor overall health and sacrificed if their body weight was 15% lower than their starting weight. Mice were killed two or five weeks after cerulein administration and pancreatic tissue collected and weighed.

### Tissue Fixation & Histology

For histological analysis, pancreatic tissue was isolated from the head and tail of the pancreas and processed as described [22]. To assess overall histology and identify differences in pancreatic tissue architecture, sections were stained with H&E. To assess fibrosis, paraffin sections were stained using Mason’s Trichrome stain (ab150686; Abcam Inc.) and fibrosis quantified using ImageJ as a percent of total tissue area. Mucin accumulation was visualized using an Alcian Blue stain kit (ab150662; Abcam Inc.) and staining quantified as a percentage of the whole tissue area.

Tissue sections were scored for ADM, PanINs and PDAC by a pathologist blinded to genotype. Progressive lesions (PanINs) were graded based on nuclear irregularities, mucinous epithelium, and dense areas of fibrosis and inflammation surrounding PanIN lesions. In all cases, 10-15 images were taken for each sample and from multiple sections at least 200 μm apart using an Aperio CS2 Digital Scanner and Aperio ImageScope software (Leica Biosystems Imaging Inc, San Diego, CA, USA) and Leica Microscope DM5500B (Leica Microsystems, Wetzlar, Germany) with LAS V4.4 software.

To assess recombination efficiency through YFP detection, tissue was fixed in 4% methanol-free paraformaldehyde for 2 hours and incubated at 4°C. Post-fixation, samples were incubated in 30% sucrose overnight at 4°C, embedded in cryomatrix (ThermoFisher Scientific), and sectioned to 6 µm using a Shandon cryostat (ThermoFisher Scientific). YFP expression was determined natively without the use of immunostaining. The percent of YFP+ cells was determined by calculating the total area positive for YFP over the total tissue area. 8-10 images per tissue were obtained with a Leica Microscope DM5500B DFC365 FX camera for analysis.

### Immunohistochemistry & Immunofluorescence

Immunohistochemistry (IHC) was performed on paraffin sections as described (Fazio et al, 2017). Following antigen retrieval, sections were permeabilized with 0.2% Triton-X in PBS, rinsed, then blocked in 5% sheep serum in PBS for 1 hour at room temperature. Primary antibodies were diluted in 5% sheep serum in PBS and incubated overnight at 4°C. Primary antibodies included rabbit α-PDX1 (1:1000; Abcam Inc. Cambridge, MA), rabbit α-amylase (1:600; Abcam Inc.), mouse α-CK19 (1:500; Abcam Inc.) and rabbit α-MIST1 (1:500; [53]. Sections were washed, then incubated in biotinylated mouse α-rabbit IgG secondary antibody (1:1000 dilution in 5% sheep serum) for 30 minutes at room temperature. Finally, sections were incubated in AB reagent for 30 minutes at room temperature and visualized using ImmPACT DAB Peroxidase (HRP) substrate (Vector Laboratories, Cat. #PK-4001/SK-4105). Slides were counterstained with hematoxylin and imaged using Leica Microscope DM5500B (Leica Microsystems) and LAS V4.4 software.

Immunofluorescent (IF) analysis was performed on paraffin embedded tissue sections similar to IHC with the exception of quenching with hydrogen peroxidase. Primary antibodies used included rabbit α-SOX9 (1:250; Millipore Sigma), rabbit α-Ki67 (1:250; Abcam Inc.) and rat α-F4/80 (1:200; Abcam Inc.). After washing, slides were incubated in α-rabbit (or rat) IgG conjugated to TRITC (1:300; Jackson ImmunoResearch, West Grove, PA) or FITC (1:300; Jackson ImmunoResearch) diluted in 5% sheep serum in PBS. Prior to mounting in Vectashield Permafluor mountant (Thermo Fisher Scientific), sections were incubated in DAPI (diluted 1:1000 in PBS; Thermo Fisher Scientific). Staining was visualized using Leica DFC365 FX camera on the Leica DM5500B microscope. Images were taken on Leica LAS V4.4 software.

### Protein Isolation & Immunoblots

For whole tissue protein, pancreatic tissue was taken from the middle portion of the pancreas and flash frozen in liquid nitrogen. Samples were processed as described (Young et al, 2018). Either 2 μg (amylase) or 40 μg of protein (SOX9, pERK and αSMA) were resolved by 12% SDS-PAGE and transferred to a PVDF membrane. Western blotting was performed as described [54]. For primary antibodies against [p] ERK (diluted 1:500) and total ERK (tERK; diluted 1:500; Cell Signaling Technology, Danvers, MA), antibodies were diluted in 0.1% TBST with 5% BSA. All other primary antibodies were diluted in 5% NFDM overnight at 4°C. Primary antibodies included rabbit α-amylase [1;1000], rabbit α-SOX9 [1:500], and rabbit αSMA [1:500]). Secondary antibody α-rabbit HRP was diluted 1:10,000 in 5% NFDM (Jackson Labs). Blots were incubated 1 hour at room temperature, washed, then incubated in Western ECL (Bio-Rad) substrate before being imaged on a VersaDoc system with Quantity One analysis software (Bio-Rad). Protein was quantified using densitometry on ImageJ and normalized to tERK accumulation.

### CyTOF – Mass Cytometry

CyTOF was carried out using Fluidigm reagents (Fluidigm Corporation; San Francisco, CA) unless otherwise noted. Whole pancreas tissue was digested with agitation in 1 mg/mL collagenase type V (Sigma) for 15 minutes at 37°C in RPMI buffer. Cell and tissue fragment mixtures were filtered sequentially through 100 and 40 micron filters and washed in ice-cold Maxpar PBS. Single cells were subjected to the Cell ID Cisplatin^ID^ reagent (1:2000 dilution) for 5 minutes at room temperature to identify live cells at the time of analysis. Samples were then stained with a panel of surface antibodies (Supplementary Table 2) for 30 minutes at room temperature according to manufacturer’s instructions. Cells were washed in cell staining buffer twice before cell fixation with 1.6% methanol-free formaldehyde (Thermo Fisher) for 10 minutes at room temperature. Samples were transferred into Nuclear Antigen Staining Buffer for 20 minutes at room temperature, then washed twice with Nuclear Antigen Staining Perm prior to intracellular staining. Intracellular antibodies were incubated with cells for 45 minutes at room temperature. Cells were then washed twice with Nuclear Antigen Staining Perm, and twice with cell staining buffer. Lastly, cells were resuspended in 1:2000 Intercalator solution in Fix and Perm buffer. Samples were acquired at the University of Rochester’s (New York, NY) CyTOF2 facility in accordance with the manufacturers protocol.

### Statistical Analysis

Data was analyzed using either a Student’s t-test (unpaired, two-tailed), one-way ANOVA or two-way ANOVA with a Tukey’s post hoc test on GraphPad Prism 6 software. Repeated measures two-way ANOVA with a Tukey’s post hoc test was used for weight loss over time. In all cases, data is shown with individual samples and error bars representing the mean ± standard error (SE). P value <0.05 was considered significant.

### Study approval

All animal experiments were performed according to regulations established by the Animal Care and Use Committee at Western University (protocol #2017-001).

## Acknowledgements

The authors wish to acknowledge the ongoing support of several national research funding agencies for this work including the Canadian Institutes of Health Research (MOP#PJT166029), the Cancer Research Society of Canada and the Rob Lutterman Foundation for Pancreatic Cancer Research. This work would not be possible without specific support from a London Regional Cancer Centre Catalyst Grant, co-supported by Mr. Keith Sammit and an internal bridge grant from the University of Western Ontario. NA and JT were funded by studentships from the Ontario Graduate Scholarships and Cancer Research and Technology Training (CaRTT) program. MK was supported by an NSERC summer studentship. JS was supported by National Cancer Institute of the National Institutes of Health under award number K08CA234222.

