## Supplemental for "Loss of Activating Transcription Factor 3 prevents KRAS-mediated pancreatic cancer"

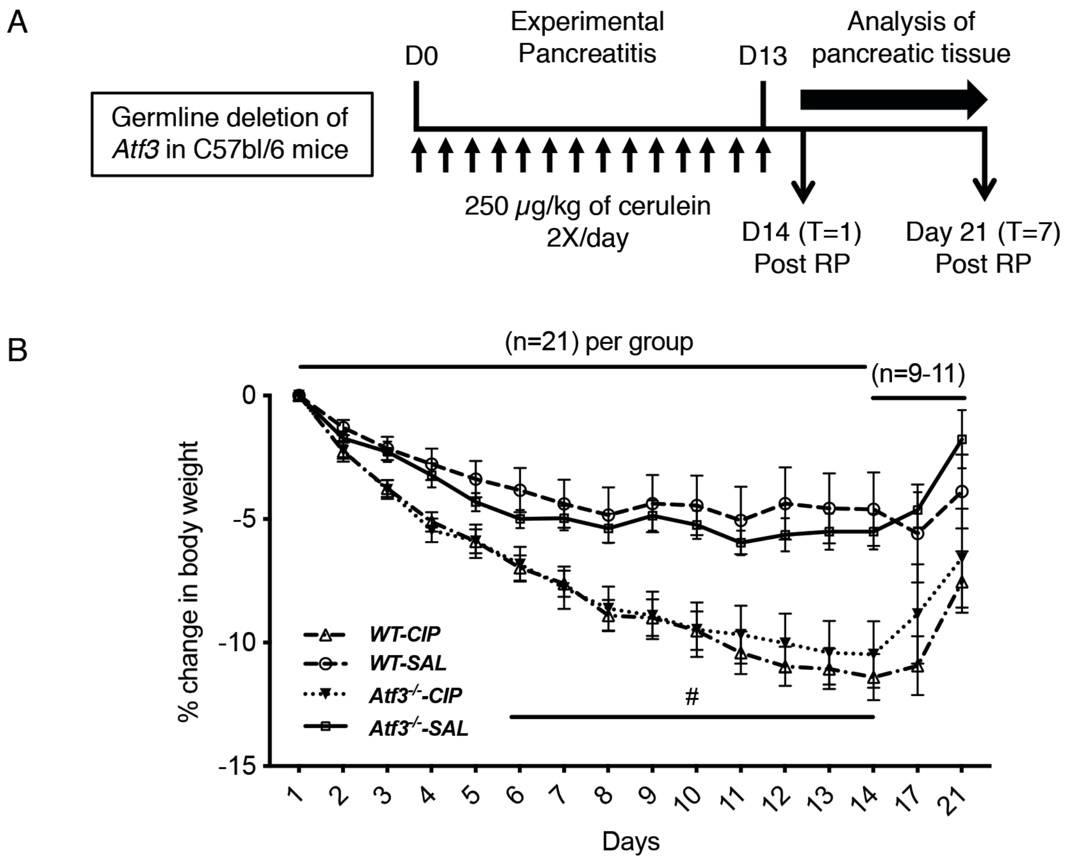


Supplemental Figure S1 (Azizi et al., 2019). *Atf3*$^{-/-}$ mice show no significant difference in body weight compared to WT mice undergoing RP. (A) A cohort of wild type (*Atf3*$^{+/-}$ and *Atf3*$^{+/+}$) and *Atf3*$^{-/-}$ mice received cerulein (250 μg/kg) or saline via intraperitoneal injections twice a day for 14 days generate recurrent pancreatic injury. (B) Body weight as a % of the starting body weight in WT and *Atf3*$^{-/-}$ mice. Mice receiving cerulein lost significant body weight compared to mice receiving saline injections (*P<0.05, #P<0.001) from days 4 to 14. However, no significant difference was observed between genotypes in the cerulein group. Error bars represent mean ± SEM. From days 1-14, n=21 for both genotypes and saline and cerulein treatment groups. After day 14, n=11 for both genotypes and treatment groups. To determine significance, a repeated measures ANOVA was performed with a Tukey' post-hoc test.

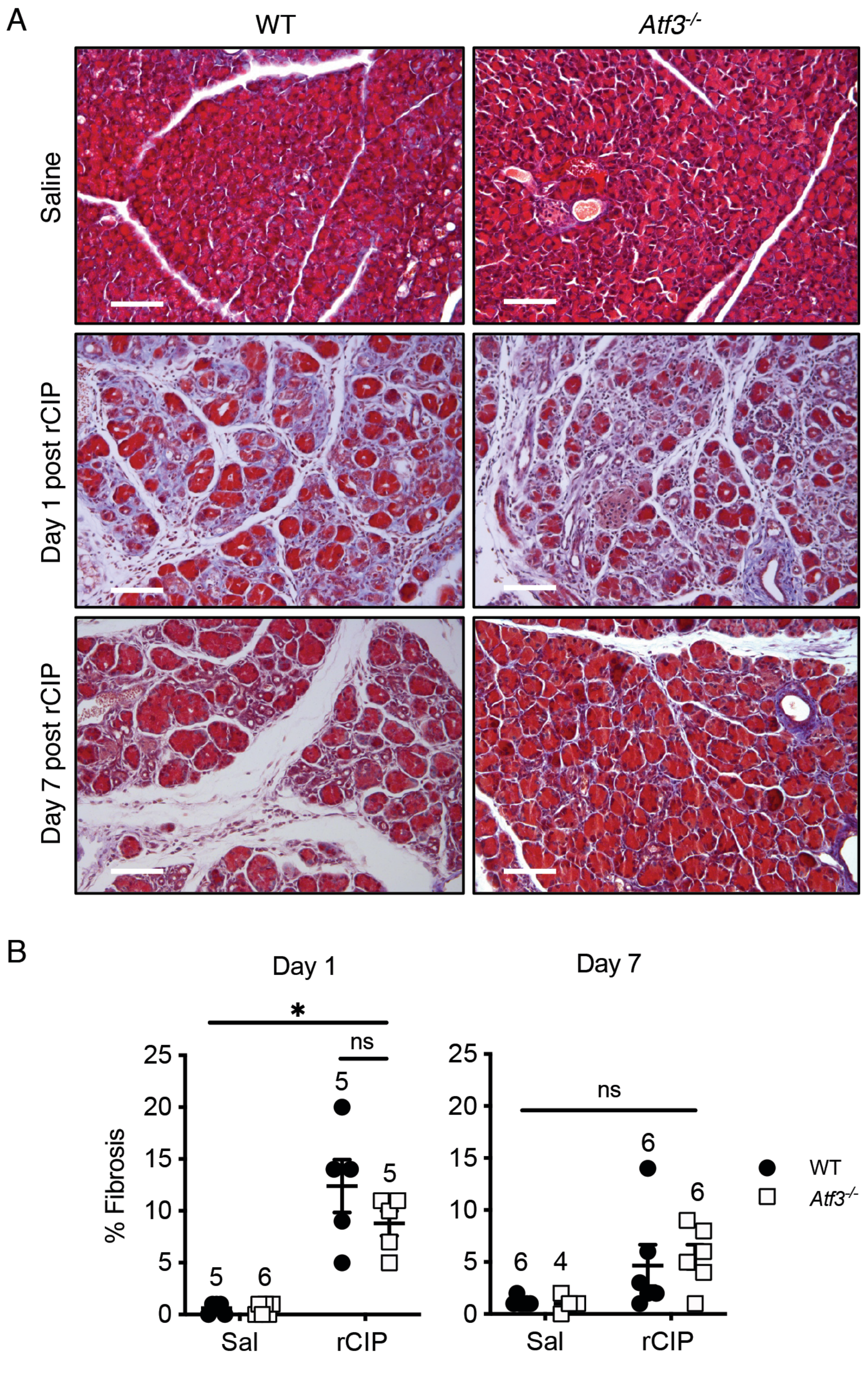


Supplemental Figure S2 (Azizi et al., 2019) The loss of ATF3 does not alter fibrosis during rCIP. (A) Representative images of Mason's trichrome staining to detect collagen as a measure of fibrosis in pancreatic tissue from saline or rCIP-treated WT and *Atf3*$^{-/-}$ CIP. Magnification bar = 140 $\mu$m. (B) Quantification of fibrosis based on green/aqua staining. *P<0.05. N values are shown above the data points. Error bars represent mean % fibrosis in tissue ± SEM. To determine significance, a two-way ANOVA was performed with a Tukey' post-hoc test.

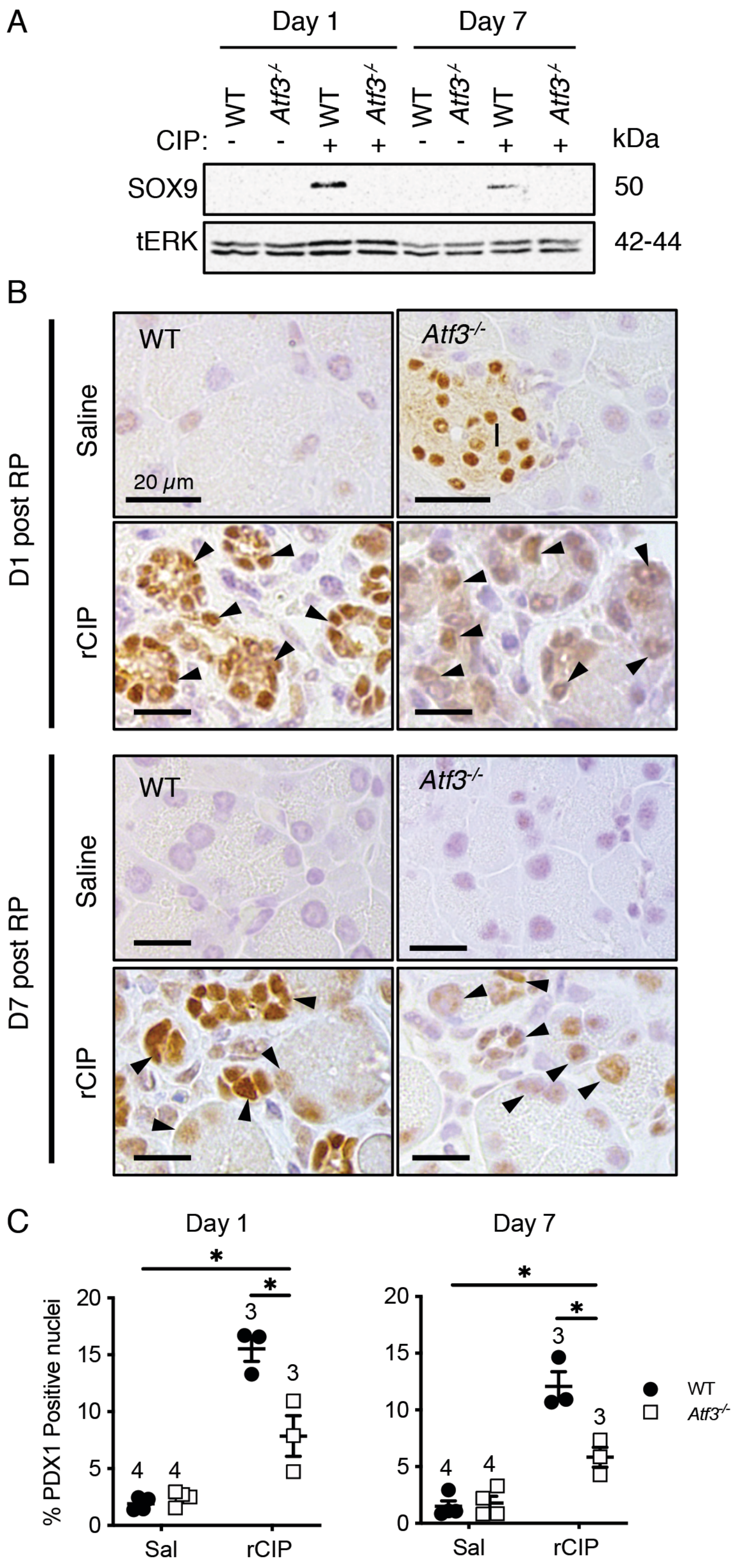


Supplementary Figure S3 (Azizi et al., 2019). *Atf3*$^{-/-}$ mice showed reduced SOX9 and PDX1 accumulation following rCIP. (A) Representative western blot analysis for SOX9 or total (t) ERK accumulation. (B) Representative IHC images show PDX1 accumulation (arrowheads) in acinar cells and putative ADM 1 and 7 days post rCIP in wild type (WT) tissue. I; islet. Magnification bar = 20 $\mu$m. (C) Quantification of PDX1+ cells from IHC images show a significant reduction in the percent of nuclei positive for PDX1 in the absence of ATF3 at both time points. N values are indicated above data points. *P<0.05; bars represent % mean ± SEM. To determine significance, a one-way ANOVA was performed with a Tukey' post-hoc test.

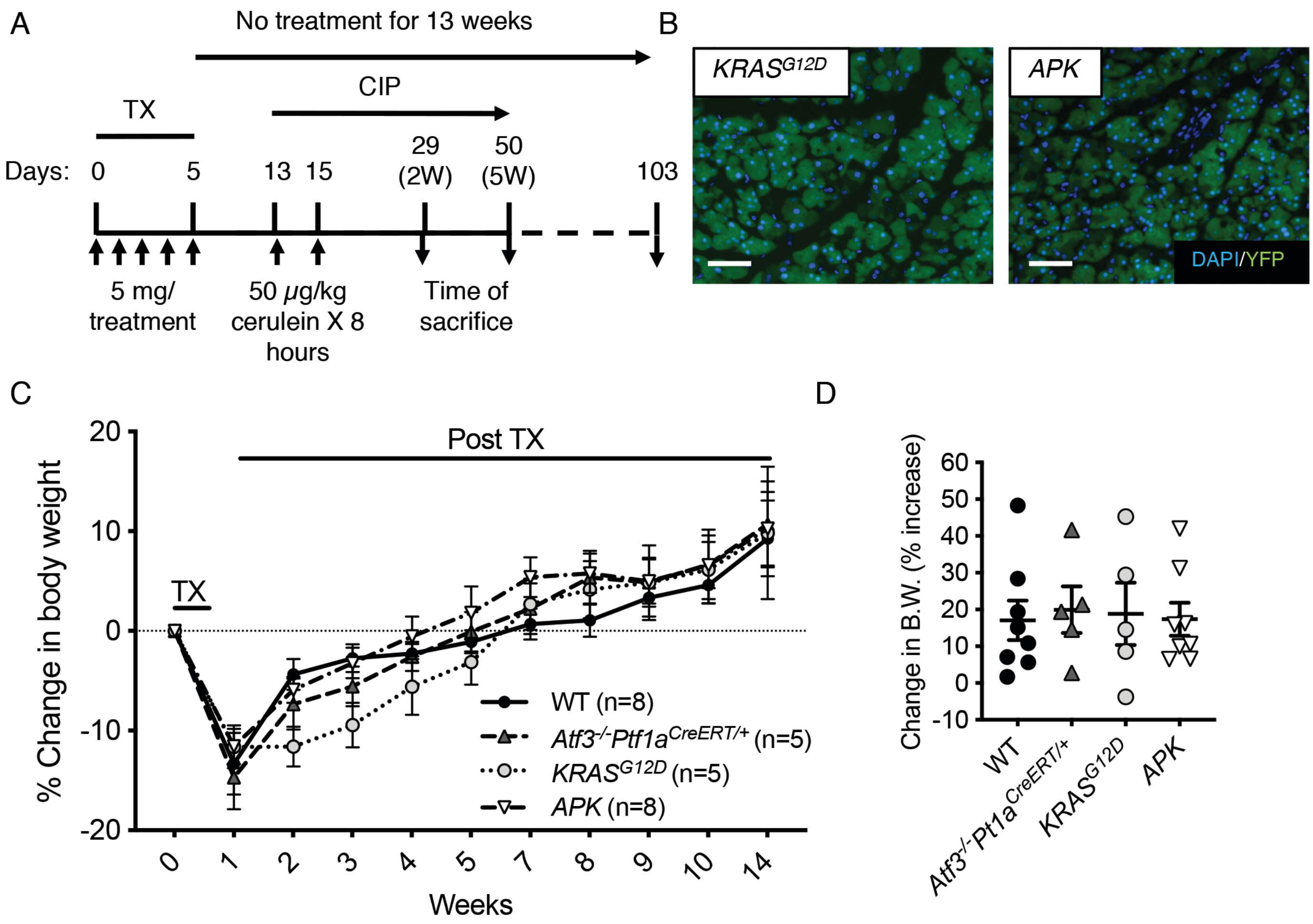


Supplemental Figure S4 (Azizi et al., 2019). Weight loss analysis of *APK* mice following activation of KRAS +/- cerulein treatment. (A) Experimental design for activating oncogenic KRAS (KRAS$^{G12D}$) +/- cerulein treatment in *Atf3$^{-/-}$* mice. Mice were sacrificed at 2 or 5 weeks (W) following cerulein treatment.
(B) Representative YFP expression following tamoxifen (Tx) induction in *Ptf1a$^{creERT/+}$Kras$^{G12D/+}$* and *APK* mice. Tissue is counterstained with DAPI to reveal nuclei. Magnification bar = 50 $\mu$m. (C) Percent change in starting body weight over the course of the experiment following activation of KRAS$^{G12D}$ with tamoxifen (TX) in 2-4 old mice. (D) Cumulative change in body weight 103 days (13 weeks) following activation of KRAS$^{G12D}$ after last Tx treatment. Percent change in body weight showed no difference between the four genotypes treated with saline. N values are indicated in brackets in C; error bars = % mean ± SEM. To determine significance, a (C) repeated measure ANOVA or (D) one-way ANOVA was performed with a Tukey' post-hoc test.

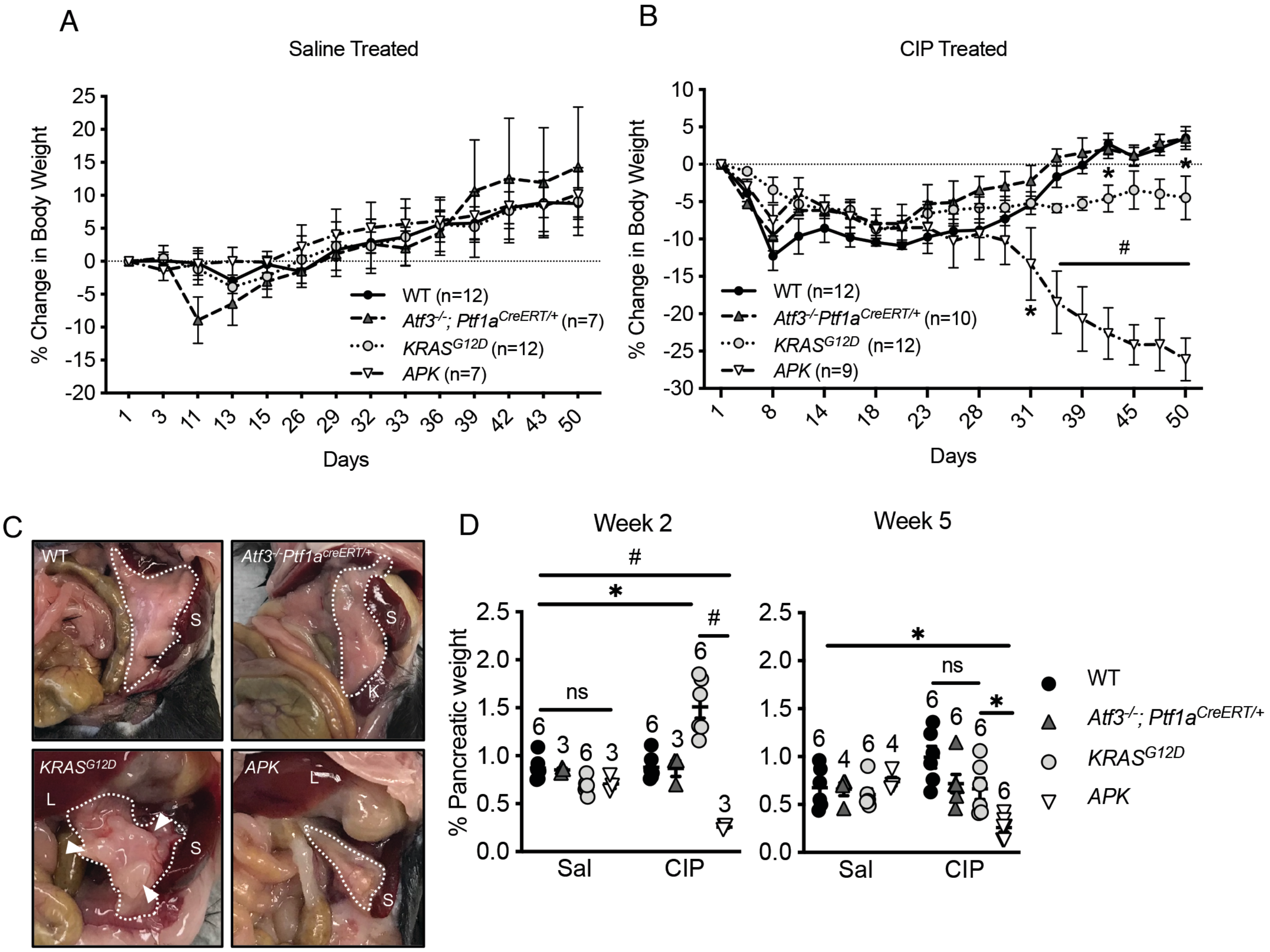


Supplementary Figure S5 (Azizi et al., 2019). Oncogenic KRAS promotes pancreatic atrophy in *Atf3*$^{-/-}$ mice. Percent change in body weight over the course of the experiment following activation of KRAS$^{G12D}$ with tamoxifen (TX) in 2-4 old mice treated with (A) saline or (B) cerulein. N values are indicated in brackets. (C) Representative images of pancreatic gross morphology (pancreas highlighted by the dotted white line; S = spleen; L = liver; K= kidney) two weeks after after CIP administration. *Ptf1a*$^{creERT/+}$*KRAS*$^{G12D}$ mice show fibrotic nodules in the pancreas (white arrowheads), while *APK* pancreata appear smaller than other genotypes. (D) Quantification of pancreatic weight as a percent relative to body weight two and five weeks post CIP. *Ptf1a*$^{creERT/+}$*KRAS*$^{G12D}$ mice show increased pancreatic weight compared to *Ptf1a*$^{creERT/+}$*KRAS*$^{G12D}$ mice treated with saline. At 5 weeks, only *APK* mice treated with cerulein had smaller pancreata compared to other groups. N values are indicated above data points; *p<0.05, #p<0.001; error bars represent the mean % pancreatic to body weight ± SEM. To determine significance, a (A, B) repeated measures or (D) two-way ANOVA was performed with a Tukey' post-hoc test.

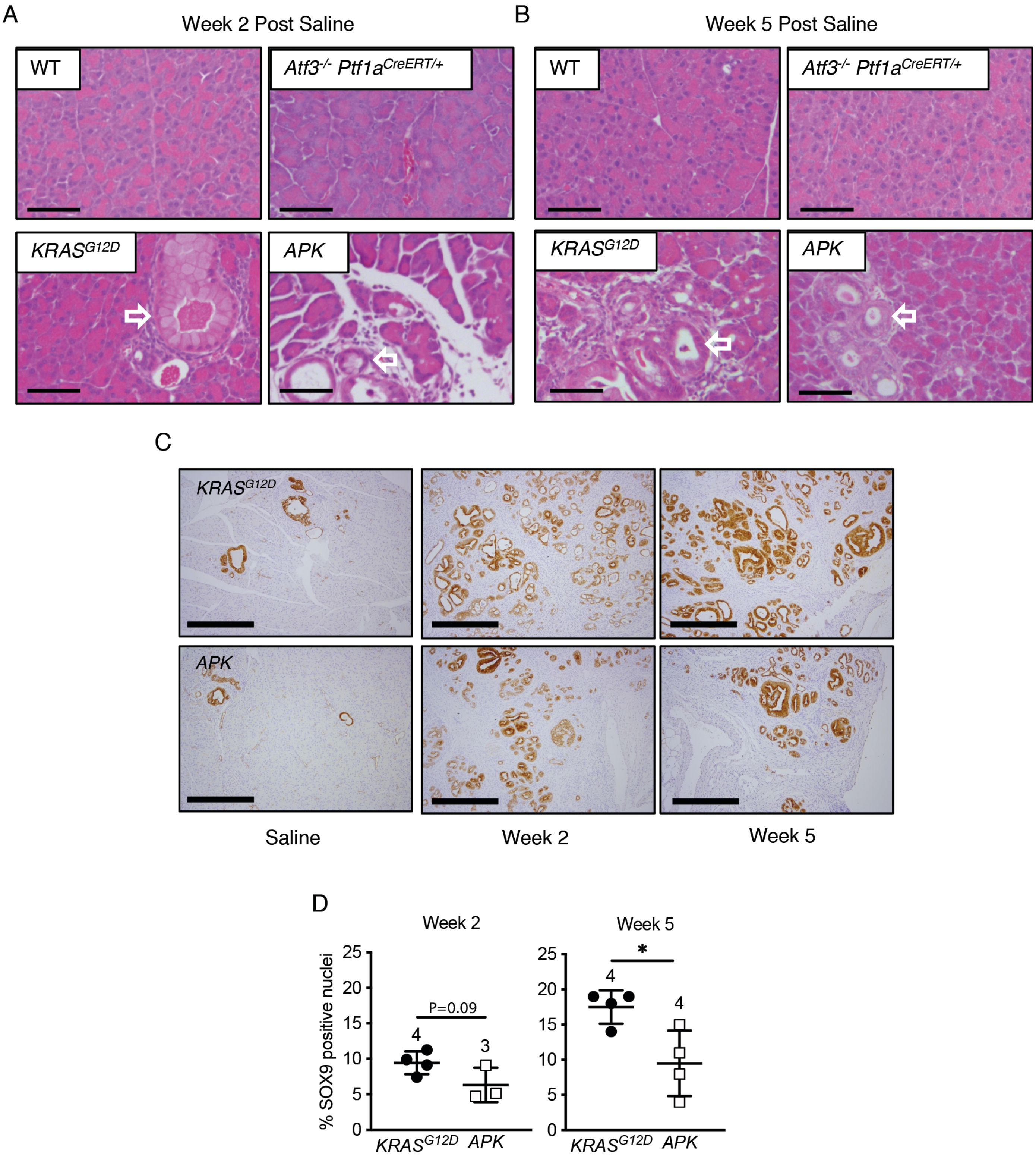


Supplementary Figure S6 (Azizi et al., 2019). Reduced PanIN progression in *APK* mice. Representative H&E analysis of mice from all genotypes two (A) and five (B) weeks after saline treatment. Arrows indicate ADM and lower grade PanIN lesions. Magnification bar = 100 $\mu$m. (C) IHC for CK20 on pancreatic sections from saline or cerulein-treated mice 2 or 5 weeks after treatment. Magnification bar = 500 $\mu$m. (D) Quantification of SOX9 staining at 2 or 5 weeks after cerulein treatment based on IF from Figure 6D. N values are indicated below graph; *p<0.05; error bars represent the mean % of SOX9+ cells ± SEM. To determine significance, a student's t-test was performed.

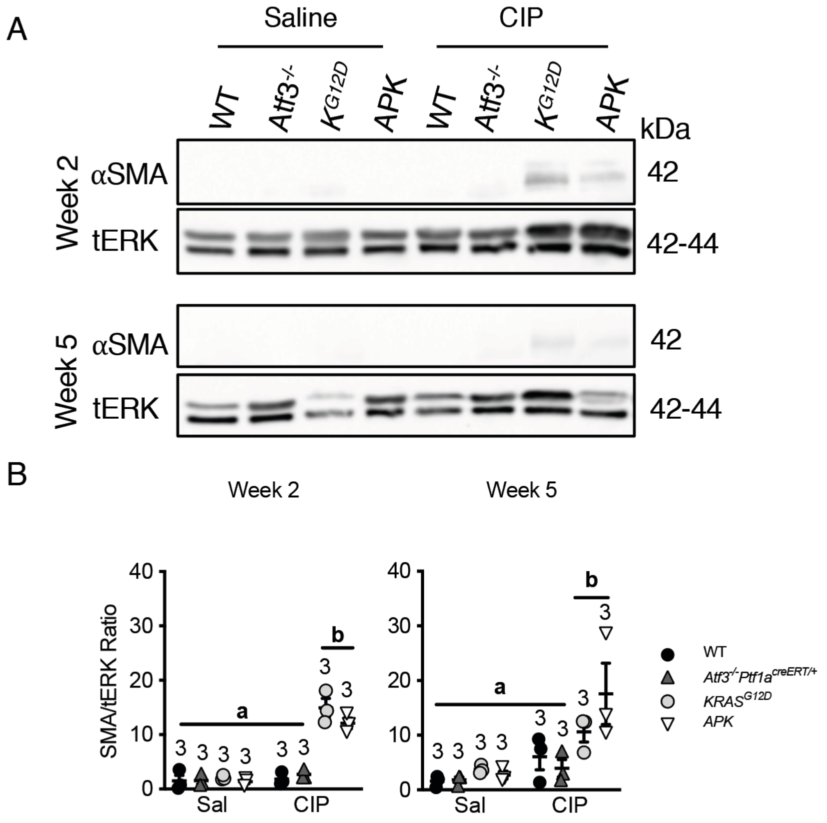


Supplementary Figure S7 (Azizi et al., 2019). Analysis of smooth muscle actin accumulation. (A) Representative western blot analysis for alpha smooth muscle actin (αSMA) and total ERK (tERK) two and and five weeks after saline or cerulein (CIP) treatment. (B) Quantification of αSMA staining normalized to tERK. N=3; letters indicate significantly different values; error bars represent the ratio of αSMA/tERK ± SEM. To determine significance, a one-way ANOVA was performed with a Tukey' post-hoc test.
